# Ontological Analysis of Brain Proteostasis Highlights the Sex-Dependent Trajectory of ApoE Isoform-Specific Regulation

**DOI:** 10.64898/2026.06.29.735293

**Authors:** Ariel E. A. Denos, Elise Clark, Noah E. Earls, Nazanin Paymard, Jared M. Elison, Benjamin S. Jones, Ethan G. Smith, Esteban G. Colman, Coleman O. Nielsen, Martin Sorensen, Noah G. Moran, Jason G. Wells, Rebecca S. Burlett, Eleni S. Vickers, Christian T. Garrard, Joshua R. Brown, Katherine L. Brown, Jossue D. Matute, Wejdene Daouahi, Mitchell F. Poulson, H. Dennis Tolley, JC Price

## Abstract

Apolipoprotein E (ApoE) is the strongest genetic predictor of Alzheimer’s disease (AD) risk, with ApoE4 increasing and ApoE2 decreasing risk relative to ApoE3. Using a global LC-MS proteomic approach, we integrated protein abundance and kinetics in Human-APOE knock-in mice for young (3-month) and aged (18-month) cohorts to quantify the changes in steady-state proteostasis. By mapping 6,052 identified proteins and 3,986 associated turnover rates into ontological groups, we observed that vesicle trafficking and mitochondrial dysregulation occur as early as 3 months in ApoE4 mice accompanied by hyperactive metabolism that eventually reduces with age. In contrast, young and old ApoE2 mice retain similar signatures to ApoE3 mice in metabolic, mitochondrial, cellular regulation, and membrane trafficking ontologies. We found that females had more isoform-induced ontological changes relative to ApoE3, providing insight into sex-dependent vulnerabilities. Our global proteomic approach for ApoE proteostasis crucially unifies independent literature observations while providing turnover kinetics to uncover the underlying mechanism behind abundance changes. Data are available via ProteomeXchange with identifier PXD079261.

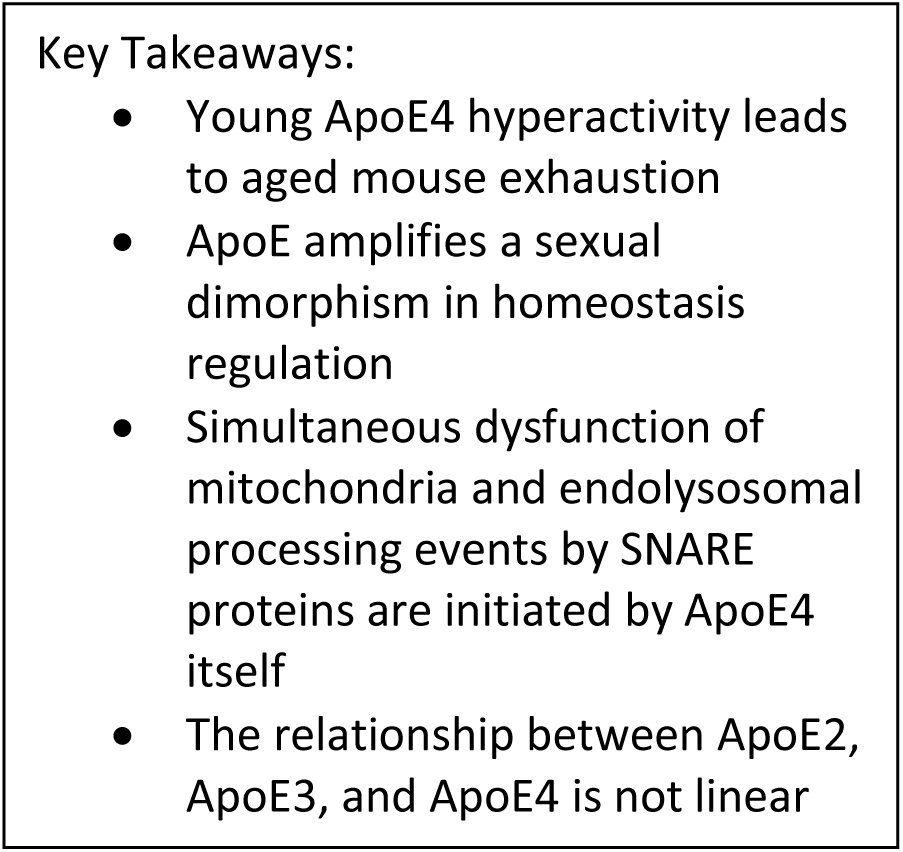

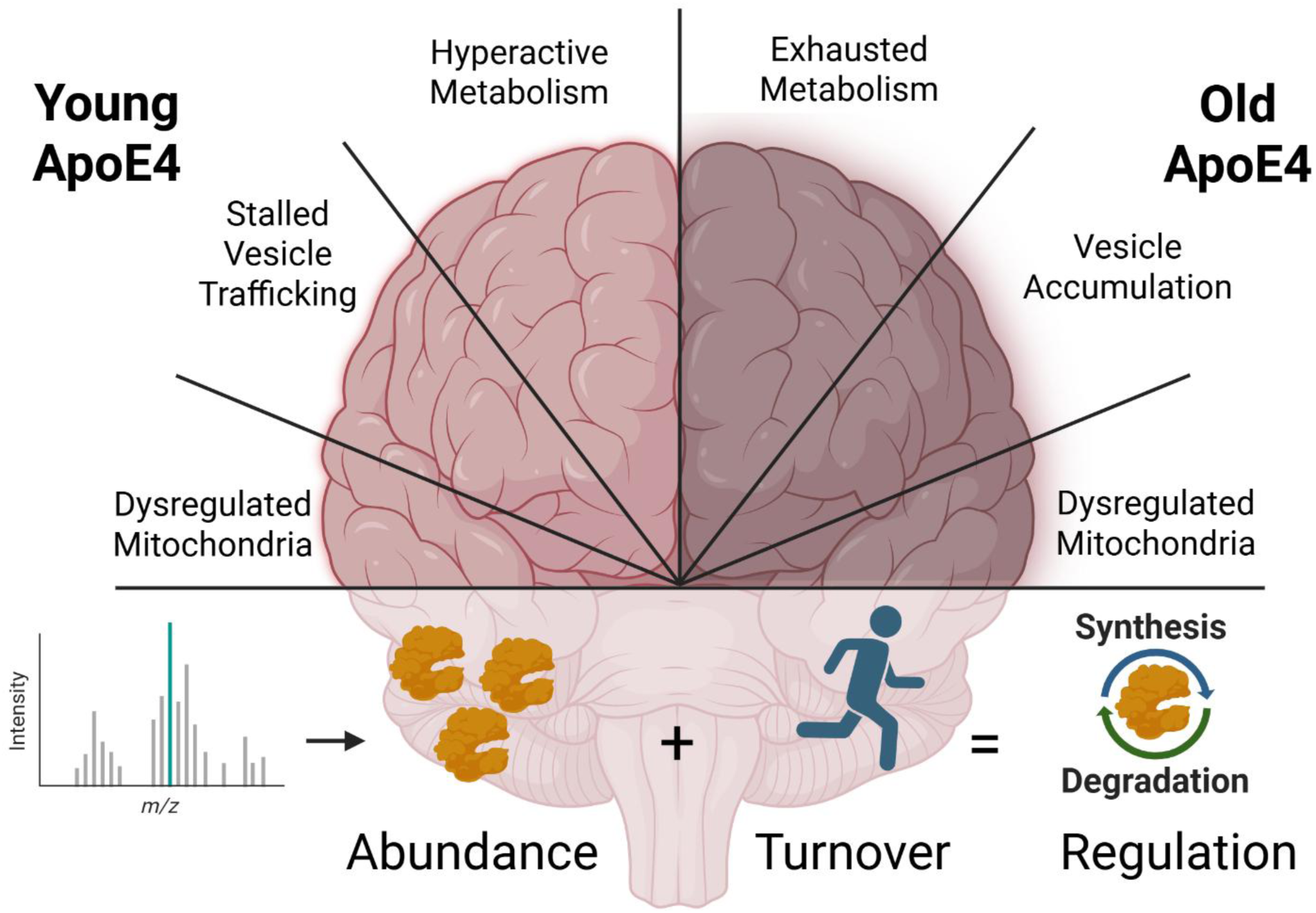

## Introduction

Apolipoprotein E (ApoE) is the primary lipid transporter in the brain and is predominantly synthesized by astrocytes, though neurons and other cell types may synthesize ApoE in stressed conditions^1^. In humans, ApoE represents the strongest genetic variant associated with Alzheimer’s Disease (AD) and exists as three common polymorphisms; ApoE2, ApoE3, and ApoE4. Although ApoE2 and ApoE4 only differ from ApoE3 by one amino acid, AD risk is heightened by ApoE4 expression and lowered by ApoE2^2,3^. In addition to the genetic component, females and the elderly are disproportionately at risk for AD development^4^. These alterations of ApoE-isoform specific risk may reflect variabilities in lipid transportation^5,6^, receptor interactions^7,8^, or downstream proteomic quality control^3,9^.

Pathologically, Alzheimer’s disease is defined by aggregation and impaired clearance of proteins^4^, suggesting a fundamental dysregulation in protein homeostasis compared to healthy individuals **(Figure 1A)**. While independent studies have linked ApoE4 to mitochondrial dysfunction^3,10^, calcium imbalances^11,12^, and stalled endocytic processing^3,13,14^, the relationship and causal-hierarchy between them remains a subject of debate^13,15–17^. Furthermore, many of these studies employ traditional biochemical techniques including western blot, immunoprecipitation, or fluorescent imaging. While informative, this provides an isolated measure of static abundance for selected features but fails to capture flux information.

**Figure 1.**
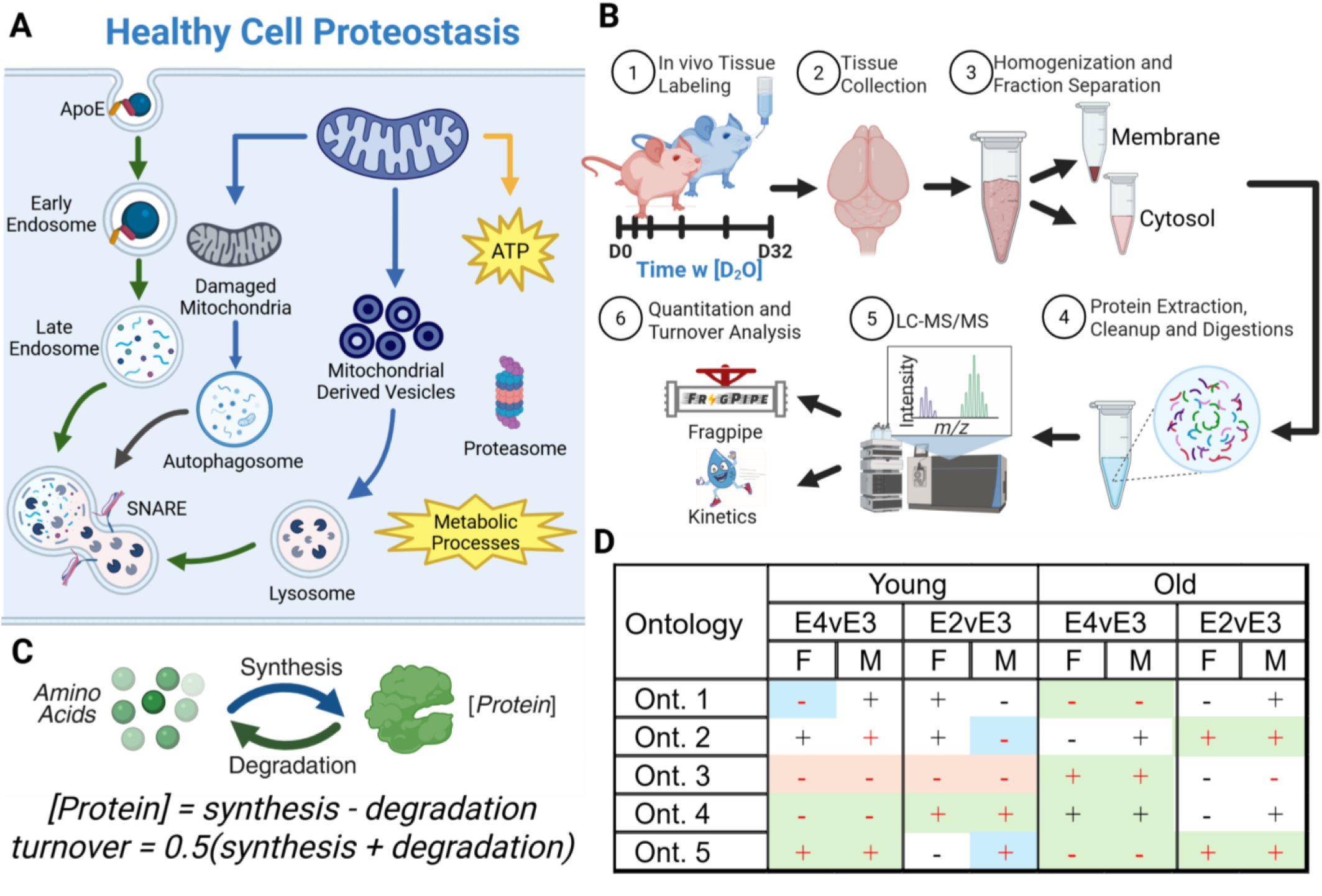
Conceptual Schematic and Experimental Outline. **(A**) Protein homeostasis schematic for healthy systems, **(B)** Male (blue) and Female (pink) ApoE2, ApoE3, and ApoE4 mice were provided with deuterium enriched drinking water. Tissues were separated into membrane and cytosolic fractions and analyzed via LC-MS. **(C)** Simplified schematic of the protein life cycle, including simplified equations for protein concentration and turnover rate. **(D)** Data interpretation schematic. Red text indicates a significant p-value and FC for an ontology, green boxes indicate a genotype specific ontology change, blue indicates early-onset ontological change, and orange indicates a significant change that is not fully interpreted as genotype specific.

Here we utilize label-free quantitative and deuterium-based kinetic proteomic measurements to quantify proteostasis fluctuations between ApoE isoforms **(Figure 1B)**. We employ a simple model to describe the relationship between protein abundance and turnover through the lens of synthesis and degradation **(Figure 1C)**. By including kinetic measurements, we uncover the underlying mechanisms that govern protein abundance changes in young and aged cohorts, and between male and female cohorts. Finally, by comparing these signatures against the proteome of ApoE2 mice, we propose a holistic mechanism of ApoE4 dysfunction and ApoE2 stability.

## Materials and Methods

### Ethics Statement

All animal handling experiments were reviewed and authorized by the Brigham Young University Institutional Animal Care and Use Committee (IACUC protocol #191102).

### Mouse Breeding

Three founding mouse lines (F_0_) were purchased from The Jackson Laboratory (JAX #029018, JAX# 029017, JAX# 027894) with heterozygous expression of human ApoE isoforms and wild-type (WT) murine ApoE (APOE2/ApoE^wt^, APOE3/ApoE^wt^, APOE4/ApoE^wt^). To reduce genetic variables and improve genomic stability of the mice, a multigenerational breeding scheme **(Figure 2)** was established to generate 3 distinct homozygous cohorts: APOE2/2, APOE3/3, APOE4/4. Genotype was verified via PCR-based amplification and agarose gel electrophoresis. Mice from breeder Cohort 1 were used to produce a generation of heterozygous APOE2/3 and APOE4/3 mice (breeder cohort 2) verified using Taqman Real Time qPCR. The final experimental cohort was the resulting homozygous APOE2/2, APOE4/4, paired with APOE3/3 littermates from breeder cohort 2 generation crosses of APOE2/3 with APOE2/3, and APOE4/3 with APOE4/3.

**Figure 2.**
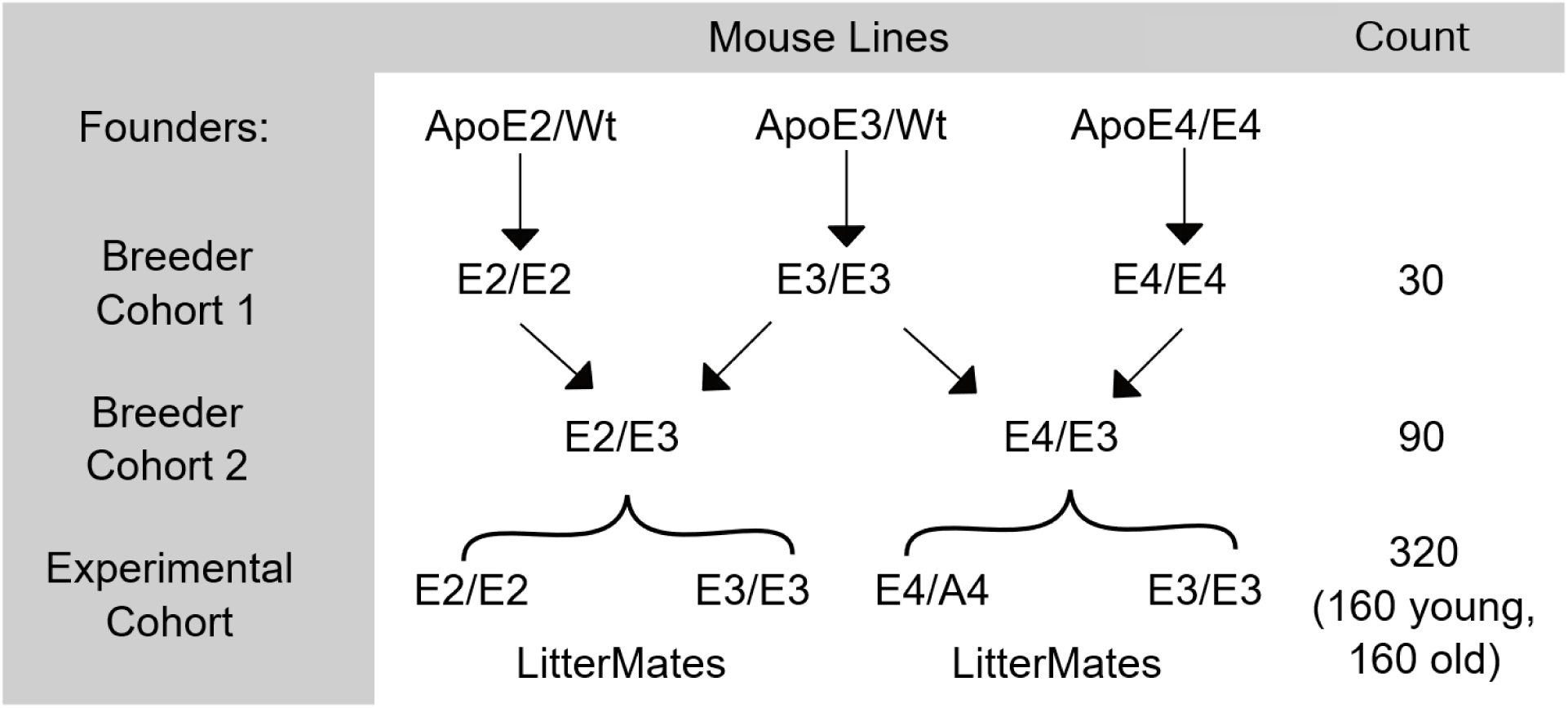
**Breeding Scheme**. Founding generations were purchased from the Jackson Laboratory and bred to produce the experimental cohort.

### Mouse Organization

This study included a total of 224 homozygous human ApoE knock-in mice, consisting of 112 young (3 months) and 112 old (18 months) mice **(Figure 3)**. An equal distribution of males and females were sacrificed for this study, with a genotype distribution of 28 ApoE2, 56 ApoE3, and 28 ApoE4 mice per age group. To minimize environmental variance, ApoE2 and ApoE4 mice were housed with ApoE3 littermates, allowing for gender and age-matched co-housed controls (cage-mates). A time-course of deuterium enrichment was established with an initial intraperitoneal injection of 0.035 ml/gram of body weight of 0.9% saline in 100% D_2_O followed by 8% D_2_O drinking water for a 32-day period. Two biological replicates with their cage mates were sacrificed at 0, 1, 2, 4, 8, 16, and 32 days post-injection. Experimental samples with cagemates were blocked by age, gender, comparison, and replicate; following tissue homogenization, samples were further divided by subcellular fraction, resulting in a final grouping of n=14 mice per sample group, with 16 sample groups per age.

**Figure 3.**
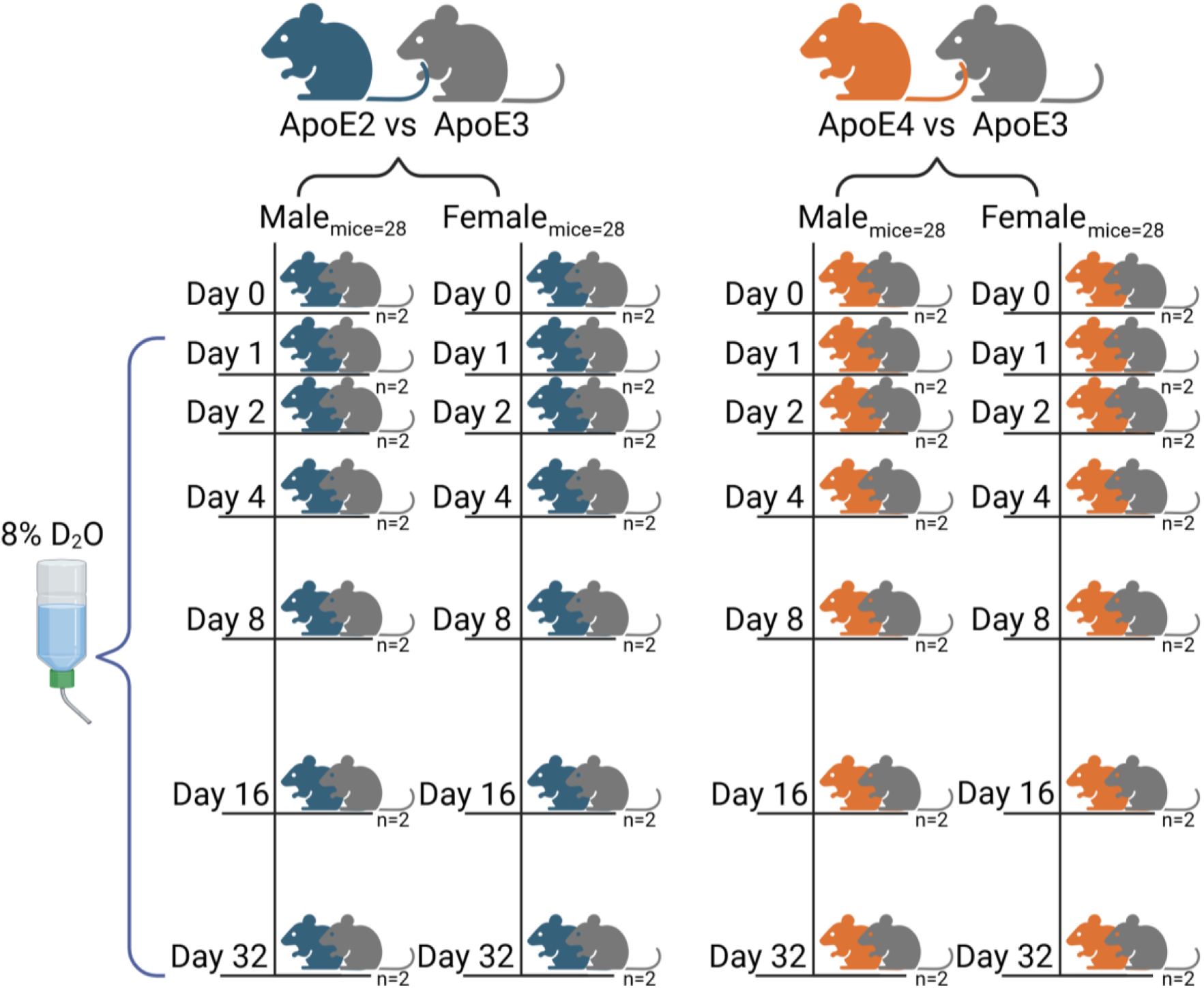
**Experimental Design**. ApoE2 and ApoE4 mice were housed with ApoE3 littermates. Male and Female mice were equally represented in the study across a 32-day time-course with 8% Deuterium Incorporation.

### Genotype Verification

All experimental mice were genotypically verified using a tail biopsy followed by DNA extraction and Taqman Real Time qPCR. Tail snips were collected at 21 days of age for genotype determination prior to organization for the experiment as well as confirmation of genotype after the organ harvest.

### Sample Collection

Mice were euthanized via CO_2_ asphyxiation followed by bilateral thoracotomy. Blood for D_2_O concentration analysis was collected by severing of the vena cava after which the resulting pool was collected from the thoracic cavity with a sterile syringe. Blood samples were placed on ice during sample collection and then spun to separate out plasma at 20,000 xg for 15 min at 4°C. To remove blood in peripheral organ samples, the mice were perfused using cold phosphate buffered saline (PBS) prior to organ collection. Brains were separated into hemispheres and the right brain was utilized for this study. Separated blood components and harvested organs were flash frozen on solid CO_2_ prior to long term storage at – 80°C. To determine deuterium enrichment for each mouse, 100 µL of red blood cell pellet was distilled in a sand bath and analyzed on a liquid water isotope analyzer (Los Gatos Research cavity ring-down spectrophotometer) as previously described^3^.

### Sample Preparation

Brain samples were homogenized following the protocol described by Zuniga et al.^3^, with the following modifications to the buffer composition and pellet processing. Briefly, tissue preparation was performed in 50mM Tris/HCl (pH to 7.55 and then treated with diethylpyrocarbonate 0.1% and ThermoScientific Halt Protease & Phosphatase Inhibitor Cocktail 0.6%). The cytosol and membrane were separated by centrifugation at 21,000 x g for 5 minutes. The membrane pellet was washed three times by resuspending in homogenization buffer then centrifugation at 21,100 x g for 5 minutes. Final fractions were resuspended in 2x lysis buffer (10% SDS, 100 mM Tris in ddH2O pH to 8.5) for subsequent analysis.

Protein concentration was measured via Pierce BCA Protein Assay Kit (ThermoFisher Scientific). For each sample, 100 µg of protein was prepared via S-Trap micro (Protifi) according to manufacturer’s recommended procedure. Modifications includ increasing the initial lysis volume to 50 µL, necessitating proportional volume adjustments of the reducing, alkylating (chloroacetamide), acidifying, and washing reagents according to the manufacturer’s *Recommendations for Various Starting Volumes.* To clean the brain samples of the lipid matrix, two optional 50% Methanol/50% Isopropanol washes were included. Samples were incubated with Trypsin Lys-C, eluted, dried via speed-vacuum, and resuspended in 3% acetonitrile and 0.1% formic acid for a final peptide concentration of 2 µg/µL.

### LCMS Analysis

All samples were run on the LC-MS/MS (ThermoFisher Orbitrap Eclipse) with a coupled Ultimate 3000 RSLC and EasySpray C18 column. Samples were analyzed multiple times with methods specific to optimize the data type. To enhance spectral accuracy and capture high fidelity isotopic envelopes for kinetic analysis, samples were analyzed in MS1 Only Acquisition. For abundance analysis, early timepoints (Day 0, 1, and 2) were run via Data-Independent Acquisition. Abundance and Kinetic acquisition methods were run as a single batch per sample group in a randomized blocked fashion to ensure retention time alignment between methods. The DIA files were processed via Fragpipe using a 1% FDR in DIA-NN quantification and outputs were used to create guide files for kinetic data by providing mass and retention time for peptide identification. DeuteRater_v7 was used for kinetic analysis^18^. Detailed methods for the LCMS analysis are available at ProteomeXchange PXD079261

### Protein Ontology Analysis

All DIA-NN LFQ outputs were normalized with the Aguilan method^19^ for each respective sample group (separated by sex, genotype comparison, fraction, and replicate) and fold change was calculated for Day 0, 1 and 2 (n=3). Replicate fold changes were then taken together to calculate the average fold change (n=6).

Statistical significance for individual proteins was determined using a heteroscedastic t-test to account for unequal variance between replication, followed by Benjamini Hochberg correction to reduce false positives. STRING (v12.0) was used with all quantified proteins as the gene set and analyzed with the default background. Ontological significance was assessed using a Wilcoxon rank-sum test based on the median fold change of all proteins within a particular ontology. Significant ontologies were those whose median exceeded a log_2_(Fold Change) of |0.07| and had a p-value less than 0.05.

### TEM Sample Preparation

For the transmission electron microscopy (TEM) sample preparation, we utilized the generic protocol from the ThermoFisher Scientific manual, *Electron microscopy in the life sciences*, with specific modifications to the fixation and infiltration stages to optimize the visualization of tissue mitochondrial architecture. Mice from the aged mouse cohort were perfused with the cold PBS and brains were harvested and placed on ice, followed by isolation of the hippocampus. Tissue was acquired and immediately trimmed into small blocks (< 0.5 mm in one axis and ≤ 2 mm in other axes) to ensure rapid chemical stabilization. Primary fixation was performed in 2% glutaraldehyde in buffer for a minimum of 2 hours, followed by five to six washes in PBS for 10 minutes each. Secondary fixation was conducted for no more than 2 hours using a 1:1 ratio of 2% potassium ferrocyanide and 2% osmium tetroxide (OsO4) in buffer, a modification designed to enhance membrane contrast for the analysis of mitochondrial complexity^20^. Following secondary fixation, samples were washed with distilled water (5–6 times for 10 minutes) and optionally subjected to in-block staining with 0.5% uranyl acetate (UA) overnight. Tissues were then dehydrated through a graded series of acetone in ethanol (10%, 30%, 50%, 70%, and 95%), finishing with three 100% acetone washes for 10 minutes each. Infiltration was achieved using Spurr’s Low Viscosity resin; samples were first placed in a vial 2/3 full of 100% acetone, filled to the top with 100% Spurr’s resin, and placed on a shaker for 1 hour. This process was repeated by exchanging the mixture for a higher ratio of Spurr’s resin, followed by a final 1-hour exchange with fresh 100% Spurr’s resin on a shaker. Finally, samples were transferred to embedding molds with pencil-written labels, pushed to the tip of the mold, and placed in a 70°C polymerization oven overnight; polymerization was verified by ensuring the resin was firm enough that a fingernail did not leave a mark.

### TEM Imaging

For the ultrastructural analysis of tissue, resin-embedded samples were sectioned to a thickness of 80 nm with a cutting speed of 0.8 mm/second using an ultramicrotome (RMC MTX), a technique consistent with established protocols for high-resolution organelle mapping^20^. Micrographs were acquired using a FEI Tecnai TF20 transmission electron microscope (TEM) operated in scanning transmission electron microscopy (STEM) mode^21^. The instrument was operated with a condenser 2 aperture of 100, a spot size of 10, and a gun length of 6. Images were captured at a resolution of 2048 with a dwell time of 10 µs per pixel. To ensure a comprehensive evaluation of the mitochondrial population ratios and structural complexity, imaging was performed across a range of scales, from low-magnification surveys at 2 µm (9kx) to high-resolution visualizations of cristae architecture at 50 nm (410kx).

### Statistical Model

#### Linear Mixed Effects Model

A linear mixed effects model was fit, modeling the combined fraction new (cfn) measurements, which are intermediate values used in rate calculation. The cfn values were utilized to add mouse cage, instrument acquisition, and individual mice as variables, which is not possible to do for turnover rates due to the nature of the rate calculation. The fitted model is as follows:

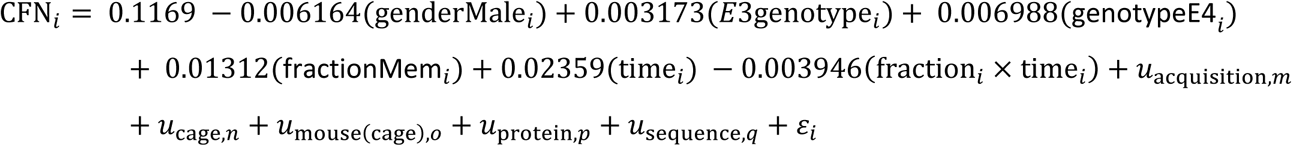

The intercept (0.1169) represents the expected cfn value for the baseline reference categories, corresponding to female mice with the E2 genotype in the cytosol fraction at Day 0 for protein A0A1W2P872 and sequence AAAAAAAAAAAAAAAASAGGK. Fixed-effect coefficients represent deviations from the intercept for each change in factor level. Random intercepts were included for instrument acquisition time, cage, mouse nested within cage, protein, and peptide sequence to account for hierarchical structure and repeated observations within the dataset. The residual term *_εi_* represents unexplained variability not captured by the terms in the model. All potential interaction terms were tested using a series of likelihood ratio tests comparing nested models.

#### Assessing Mitochondrial Structure

Images of 130 mitochondria (consisting of 74 ApoE2 mitochondria and 49 ApoE4 mitochondria) were randomly labeled and independently reviewed by a panel of 6 blinded individuals. Structural damage was assessed according to a indexed rubric of images in a 4-tiered scoring system similar to Shults^22^ and Rybka et al^23^: Type 1 (healthy/functional): Organized cristae and distinct double membrane, Type 2 (early damage): Loss of dense and organized cristae, membrane still intact. Type 3 (advanced damage): Cristae fragmentation and homogenization and possible discontinuous double membrane. Type 4 (dysfunctional): Major cristae loss and possible mitochondrial rupture. The average score of each mitochondria was calculated and used as input for a Welch’s *t*-test to determine structural variability between ApoE2 and ApoE4 genotypes.

### Kinetic Analysis

The rationale for our kinetic analysis is explained in detail by Zuniga et. al.^3^. In short, our protein homeostasis model assumes steady state kinetics with a large pool of free amino acids. That pool consists of imported and metabolized amino acids, as well as those resulting from protein degradation. Though there are several parameters accounting for the loss and accumulation of proteins and amino acids, our current analysis uses a simplified model that focuses on dominating forces. Synthesis is assumed to be the rate limiting step for protein generation, much slower than folding or t-RNA-charging with no import/export of protein (Fig 1C). As stated previously, we assume a large pool of amino acids that is well-mixed, assuming amino-acid selection is random and non-biased. Additionally, we assume the protein pool is well mixed and is subject to random protein degradation. Consistent with literature assumptions^3,24–26^, we arrive at equation 1 where a change in protein concentration is represented by the difference in synthesis and degradation rates.

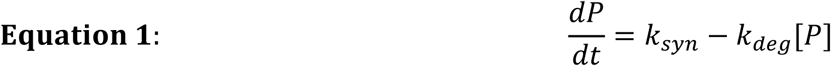

Our model utilizes a zero-order synthesis rate (k_syn_) and a concentration-dependent degradation step (k_deg_). We understand assuming first order kinetics is not a catch-all method and does not accurately describe reversible aggregation^24,27,28^, multistage synthesis rates^29–31^ and selective degradation^3,32–35^. A model accurately accounting for these variables is not currently available, and thus we, as well as others, have continued to adopt synthesis as zero-order^24–26,36^. This assumes synthesis is independent of amino acid availability and protein concentration, while degradation is only regulated via protein concentration.

Importantly, since our mice are not subject to any treatments besides deuterated water, we assume they are each in a state of undisturbed protein homeostasis specific to each isoform. This suggests that there is no reason for a spontaneous up- or down-regulation of synthesis or degradation, and assumes synthesis and degradation are equivalent. Therefore, the change in protein concentration over time is equal to 0 within a given polymorphism. However, we expect the fold-change relationship between isoform rates to be quantifiable, as described in Equation 2.

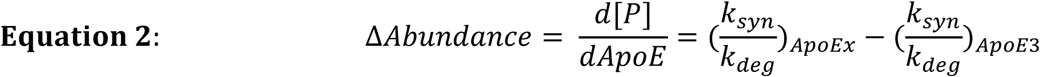

By incorporating deuterium after t=0, we can begin to measure the replacement of old unlabeled proteins [P] with new labeled proteins [P^D^]. Though [P]+[P^D^] is independent of time and will always equal k_syn_/k_deg_, the fraction of deuterated protein over time is expressed in Equation 3.

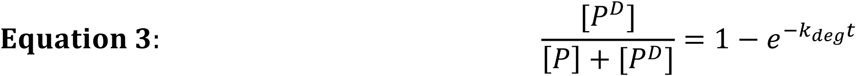

Although degradation appears to be a driving force for protein replacement, our assumptions allow us to use k_syn_/[P] in place of k_deg_. Because per-molar synthesis and degradation rates can be used interchangeably, we find it conceptually superior to define turnover rate and the mean of both rates:

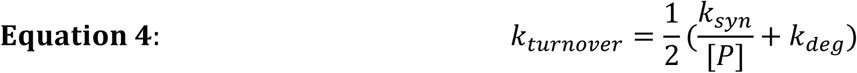

Using the simplified relationships proposed in **Figure 1C** **(Equation 5** **and 6)**, we can determine the type of regulation governing abundance changes. For instance, a relative increase in protein concentration may be attributed to either increased synthesis or decreased degradation **(Figure 4D)**. We can then deduce which type of regulation is occurring in the tissue by incorporating the turnover rate. If the turnover rate fold-change is negative, the regulation can be determined as decreased degradation. Conversely, if the turnover rate fold-change was positive, the driving mechanism for the accumulation of proteins is increased synthesis **(Figure 4D)**.

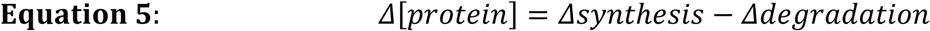

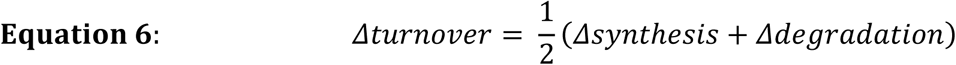

**Figure 4.**
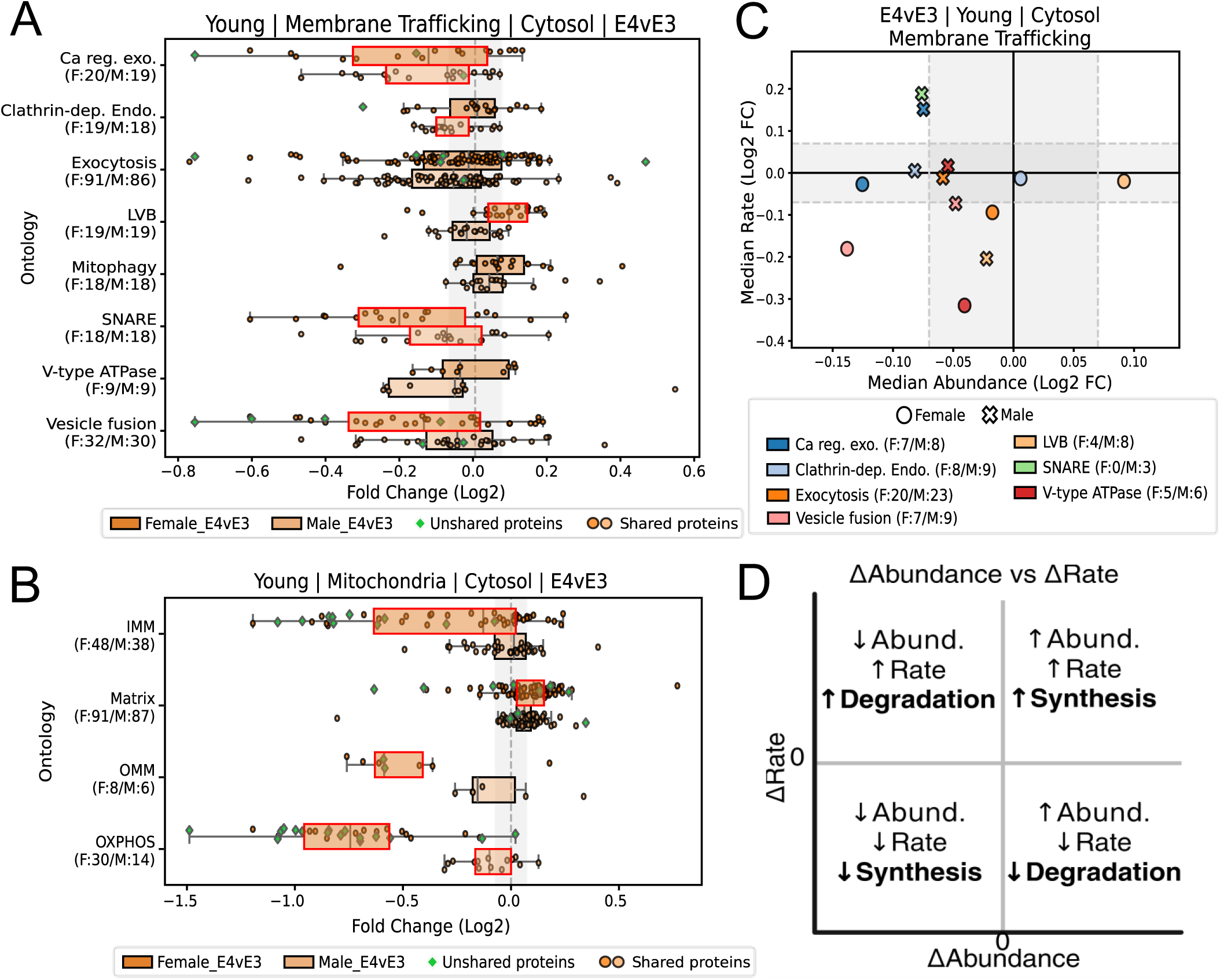
**Three-Month-Old Healthy Apoe4 Mice Exhibit Mitochondrial and Membrane Trafficking Disparities Compared to ApoE3**. Boxplots of normalized protein abundances of 3-month-old mice organized into functional groups for female (top, dark) and male (bottom, light) in **(A)** Membrane Trafficking Ontologies, **(B)** and Mitochondrial Ontologies. **(C)** Proteostasis Plot of median log2FC abundances (x-axis) and rate (y-axis) for Membrane Trafficking ontologies that exceeded the fold-change cutoff and had p-value significance. For **(A)** and **(B)**, X-axis for is log2 fold change of ApoE4/ApoE3, y-axis for A and B are ontologies. Fold Change cutoffs for abundance and rate ontologies are |0.07|. Red outlines on boxes represent a significant Wilcoxon p-value of less than 0.05. Ca reg. exo. = Calcium-ion regulated exocytosis, Clathrin-dep. Endo. = Clathrin-dependent endocytosis, LVB = Lysosome Vesicle Biogenesis, Mitophagy = Autophagy of Mitochondrion, IMM = Inner Mitochondrial Membrane, Matrix= Mitochondrial matrix, OMM = Outer Mitochondrial Membrane, OXPHOS= Oxidative Phosphorylation.

## Results

### Data Interpretation Model

The experimental genotypes (E2 or E4) were housed with E3 littermates for the entire experiment to minimize environmental variability **(see** **Figure 3).** Data were collected and are expressed as the relative change of the experimental mouse to its co-housed ApoE3 littermate control; thus results are labeled as E2vE3 or E4vE3.

Multiple studies using an ApoE Knock-in model have reported minimal pathology or cognitive decline, as reviewed by Van Heuvelen et al.^37^. Therefore, any significant deviation from ApoE3 indicates a loss of protein turnover that is sustainable, but implies a bias in proteome homeostasis that eventually leads to loss of equilibrium between biological processes^12,38^. Consequently, while the subtle proteostasis fluctuations observed in this controlled study may be well tolerated in otherwise healthy mice, they may reflect loss of proteomic equilibrium that becomes increasingly consequential in humans, where these changes reduce resilience against the impacts of aging, poor diet, environmental exposure, and physiological stressors.

We applied a multi-step interpretative framework on our ontologically categorized protein data **(Figure 1D)**. First, we evaluated the median ontology fold-change and Wilcoxon p-value, both of which had to exceed significance thresholds to be determined significant (p<0.05 and a log2(fold change) beyond |-0.07|). Second, we identified conserved proteomic signatures across both sexes within a genotype comparison. Third, where young mice exhibited unique ontological shifts not yet present in both sexes, we assessed whether these signatures emerged in both males and females in the aged mice. We propose these to be early-onset changes and represent an accelerated pathological trajectory. These early-onset signatures were particularly observed in female mice^39^, reflecting the heightened vulnerability of the female brain to ApoE isoform-mediated proteostatic stress^40,41^. Lastly, if an ontological signature was equivalent between both sexes and genotype comparisons (E2vE3 and E4vE3), it was noted as significant but not considered a driving factor for ApoE4-dependent increased AD risk. Ultimately, this framework allows for unbiased ontological selection and provides discernment between primary driving factors and secondary downstream effects.

### Proteostasis Changes are Amplified in the Cytosolic Fraction

To determine whether regulatory patterns differ between subcellular compartments, we separated brain homogenates into cytosolic and membrane fractions. Importantly, most significant changes occurred in the cytosolic fraction. To determine if this is due to heightened stability or increased variability of the membrane fraction, we performed a Levene’s test to compare the variability in abundance within the cytosolic and membrane fractions. We found a significant difference (p < 0.05) between the two fractions, with the cytosolic fraction exhibiting a variance of 0.151 compared with 0.083 in the membrane fraction. We also fit a linear mixed-effects model on the rate data to compare multiple variables in the entire dataset, and cellular fraction emerged as a significant factor with the membrane fraction exhibiting generally slower turnover rates.

Together, these data suggest greater overall proteomic stability and lower ApoE-dependent variation in the membrane fraction compared to the cytosol. By separating the fractions, we have increased the contrast of membrane and cytosol frameworks in the tissue, allowing for the distinct regulatory patterns to be elucidated. Surprisingly, while the membrane fractions between genotypes were very similar, we noted many membrane-associated proteins to be significantly altered in the cytosolic fraction. This suggests that transport of such proteins to the membrane fraction, regulation of post-translational modifications, strength of membrane associations, vesicle size, or trafficking may all be impacted by ApoE isoforms. Due to the stable composition and high-similarity of the membrane fractions across all the experimental groups, the following results and discussion section are focused on the cytosolic fraction.

### Housing Effects

Shared housing modifies gut health and microbiome which is closely related to brain health^42,43^. To assess whether a housing effect existed, we compared the two E3 control groups which were housed with the E2 or E4 experimental mice. Since isotope labeling is internally normalized the turnover rates are less impacted by run-to-run bias, our analysis focused on turnover rates. First, we calculated fold changes in turnover rates between the ApoE3 groups and performed ontology enrichment analysis followed by Benjamini-Hochberg correction; however, no ontologies had significantly changed rates. Next, we performed a two-sample t-test on the rate files by two methods: per protein and per group. The first t-test was implemented at the protein level, comparing protein-specific turnover rates between E3 mice housed with E2 mice and E3 mice that were house with E4 mice. For the second t-test, observations were averaged within each experimental group, where a group was defined by a unique combination of age, sex, cellular fraction, genotype pair, and genotype (e.g., OMBC_4v3_E3). These group level averages were then used as the response in this second two-sample t-test. The results indicate that the two E3 groups are not statistically different from each other (p=0.6981), confirming the E3 mouse brain proteome was not significantly affected by their experimental cagemate.

Lastly, we fit a linear mixed-effects model on the precursor rate file to evaluate the impact of housing while accounting for other potential sources of variation including fraction, sex, mouse-to-mouse variability, and cage effects. Consistent with the t-test results, housing does not have a significant association with the global turnover rate. Therefore, our analysis is a genuine comparison of ApoE isoform-specific proteostasis achieved with minimal interference due to microbiome effects.

### ApoE4 Drives a Simultaneous Dysfunction of Mitochondrial and Vesicle Trafficking Machinery

Mitochondrial dysfunction and disruption of endocytic vesicle processing are frequently implicated in ApoE4-associated Alzheimer’s disease risk^13,17^. Though the ApoE4 knock-in model does not develop amyloid plaques it is a model for the earliest stages of dysfunction that leads to the disease. In the cytosol of healthy young 3-month-old ApoE4 mice, we observed significant changes in both mitochondrial and vesicle trafficking ontologies across male and female cohorts. Specifically*, SNARE* protein complexes and *calcium-ion regulated exocytosis* were significantly downregulated in membrane trafficking ontologies **(Figure 4A)**. Cell-wide vesicle movement and processing requires SNARE proteins, which mediate the tethering and fusing of vesicles. In mitochondrial ontologies, we observed a reduction in *(Oxidative phosphorylation) OXPHOS* in ApoE4 mice.

Females additionally experienced ontological changes in the *Inner* and *Outer mitochondrial membranes (IMM/OMM)*, and the *Mitochondrial matrix* **(Figure 4B)**. Although many of the proteins within these ontologies are membrane-associated, these changes were only seen in the cytosolic fraction with no significant changes in the membrane fraction relative to ApoE3 **(Supplementary Figure 1A-B).** Together, these data suggest that the membrane and cytosolic fractions maintain distinct protein homeostatic states and that transport between them may be disrupted.

To uncover the underlying regulation responsible for the observed abundance changes, we integrated abundance (x-axis) and turnover rate (y-axis) on a proteostasis plot **(Figure 4D).** This allows us to determine, for example, whether a negative abundance change results from increased degradation (when turnover is increased) or decreased synthesis (when turnover is decreased). At 3 months of age, no mitochondrial ontologies had significant rates in the cytosolic or membrane fraction for E4vE3 **(Supplementary Figure 2-3)**. Intriguingly, we observed a sex-divergent regulatory response affecting related vesicular trafficking pathways: males exhibited increased degradation for proteins involved in *calcium-ion regulated exocytosis*, whereas females displayed reduced synthesis of *vesicle fusion* proteins **(Figure 4C)**.

### Metabolic Rewiring Promotes Hyperactivity in Young ApoE4 Cohorts

Given OXPHOS disruptions, we suspected additional metabolic pathways may be similarly altered. To test this, we compared the metabolic pathways provided by STRING from our protein list. We found *Glutathione metabolism* and *glycolysis* protein abundances to be significantly increased in ApoE4 mice **(Figure 5A)**. In female mice, we additionally observed increased TCA cycle and amino acid metabolism. However, *Mitochondrial Fatty Acid Beta Oxidation* was not significantly changed in ApoE4 mice, suggesting that not all mitochondrial metabolic pathways are affected. The observed increase in *glutathione metabolic* proteins for both sexes was driven primarily by increased synthesis **(Figure 5C).** On the other hand, glycolysis shows decreased degradation, indicating varying regulation of metabolic pathways **(Figure 5C)**. No metabolic ontologies were significantly changed in the membrane fraction **(Supplementary Figure 1C** **and 3)**.

**Figure 5.**
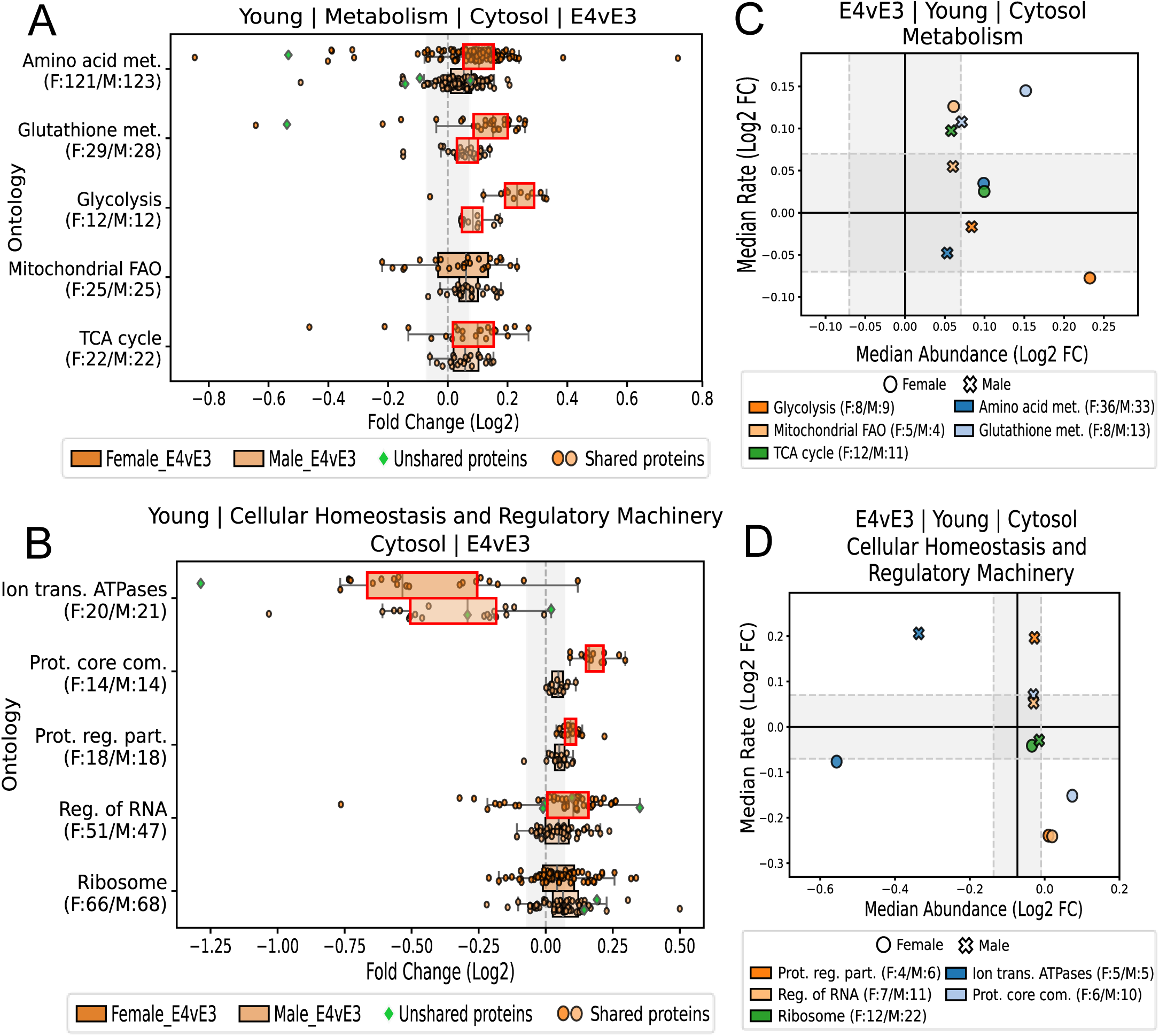
ApoE4 Mice Exhibit Increased Metabolism. Boxplots of normalized protein abundances of 3-month-old mice organized into functional groups for female (top, dark) and male (bottom, light) in **(A)** Metabolism, **(B)** and Cellular Homeostasis and Regulatory Machinery Ontologies. Proteostasis Plot of median log2FC abundances (x-axis) and rate (y-axis) for **(C)** Metabolism and **(D)** Cellular Homeostasis and Regulatory Machinery ontologies that exceeded the fold-change cutoff and had p-value significance. For **(A)** and **(B)**, X-axis for is log2 fold change of ApoE4/ApoE3 and the y-axis are ontologies. Fold Change cutoffs for abundance and rate ontologies are |0.07|. Red outlines on boxes represent a significant Wilcoxon p-value of less than 0.05. Amino acid met. = Amino acid metabolic process, Glutathione met. = Glutathione metabolism, Glycolysis = Canonical Glycolysis, Mitochondrial FAO = Mitochondrial fatty acid beta-oxidation, TCA cycle = Tricarboxylic acid cycle, Ion trans. ATPases = Ion transport by P-type ATPases, Prot. Core com. = Proteasome core complex, Prot. Reg. part. = Proteasome regulatory particle, Reg. of RNA = Regulation of RNA splicing.

We next assessed whether the proteostasis regulatory machinery was also altered. We specifically interrogated the abundance and kinetics of proteins involved in the *Regulation of RNA splicing*, the *Ribosome*, the *Proteasome,* and *Ion transport by P-type ATPases* **(Figure 5B)**. We observed an upregulation of proteins involved in the *Proteasome* and *RNA regulation* in the cytosolic fraction of females; this accumulation was driven by decreased degradation **(Figure 5D)**. Notably, *Regulation of RNA splicing* was downregulated in the membrane fraction **(Supplementary Figure 1D)**, along with the *Ribosome* complex. This may be indicative of protein translocation between the two fractions in female ApoE4 mice; where proteins in one ontology are increased in the cytosol and decreased in the membrane. Although significantly downregulated in ApoE4 mice, *Ion transport by P-type ATPases* was equally altered in ApoE2 mice relative to ApoE3 **(Figure 8D**, discussed below), therefore this ontological change is not considered a driving factor of ApoE4-induced proteostasis.

However, regulation of *ion transport by P-type ATPases* differed between ApoE4 males and females, indicating a divergent kinetic response. A standard linear regression model on protein turnover rates across the ontologies presented in the study found sex to be significant variable, with males exhibiting lower rates than female mice. These provide additional evidence of the sex-dependent regulatory disparity noted previously.

### ApoE4 Induced Hyperactivity leads to Aged Metabolic and Regulatory Exhaustion

In aged ApoE4 mice, we observed a broader reduction of metabolic ontologies. Contrary to their upregulation in young mice, *Glycolysis* and *amino acid metabolism* are significantly reduced in aged ApoE4 mice, alongside *fatty acid beta-oxidation* **(Figure 6A)**. The male protein turnover reveals that the reduction of *TCA cycle* and *fatty acid beta-oxidation* proteins is driven by increased degradation **(Figure 6C)**. However, the proteasome core complex itself is reduced in male mice **(Figure 6B),** potentially indicating reliance on non-proteasomal degradation machinery. Aged females also experience a reduction of the proteasome, though kinetic regulation of the *proteasome core* and *regulatory particle* exhibit distinct, non-parallel rate signatures **(Figure 6D)**. Of note, this proteasomal kinetic decoupling is age-specific, as the regulatory particle and core complex kinetics in young females were equivalent.

**Figure 6.**
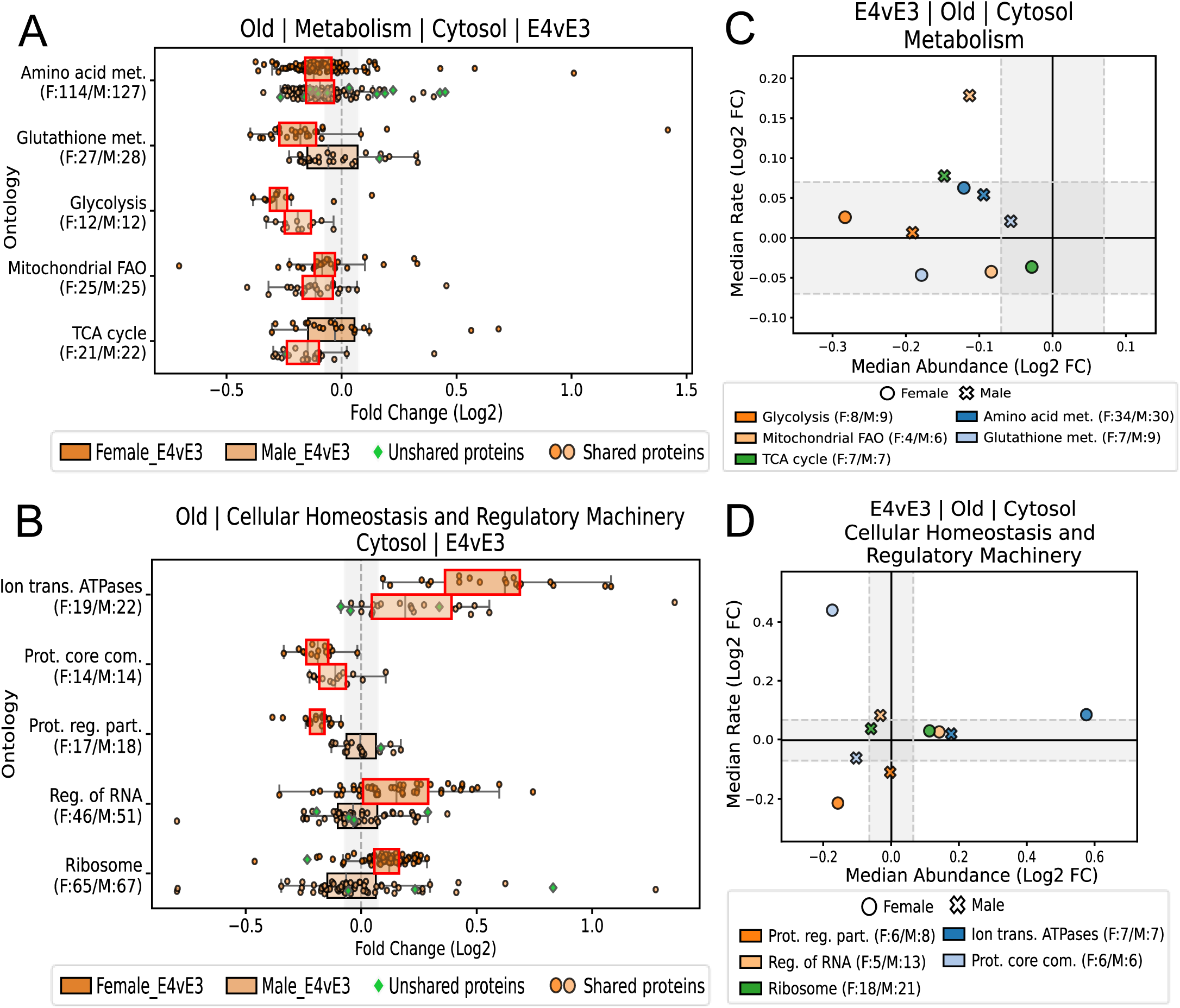
Aged ApoE4 Exhibit Reduced Metabolic and Proteosome Machinery. Boxplots of normalized protein abundances of 18-month-old mice organized into functional groups for female (top, dark) and male (bottom, light) in **(A)** Cellular Homeostasis and Regulatory Machinery, **(B)** and Metabolism Ontologies. Proteostasis Plot of median log2FC abundances (x-axis) and rate (y-axis) for **(C)** Cellular Homeostasis and Regulatory Machinery and **(D)** Metabolism ontologies that exceeded the fold-change cutoff and had p-value significance. For **(A)** and **(B)**, X-axis for is log2 fold change of ApoE4/ApoE3 and the y-axis are ontologies. Fold Change cutoffs for abundance and rate ontologies are |0.07|. Red outlines on boxes represent a significant Wilcoxon p-value of less than 0.05. Amino acid met. = Amino acid metabolic process, Glutathione met. = Glutathione metabolism, Glycolysis = Canonical Glycolysis, Mitochondrial FAO = Mitochondrial fatty acid beta-oxidation, TCA cycle = Tricarboxylic acid cycle, Ion trans. ATPases = Ion transport by P-type ATPases, Prot. Core com. = Proteasome core complex, Prot. Reg. part. = Proteasome regulatory particle, Reg. of RNA = Regulation of RNA splicing.

Of other protein regulatory ontologies, *regulation of RNA splicing* in females maintains a positive abundance as seen in young mice **(Figure 6B)** coupled with increased *ribosomal* proteins. This is a unique directional preservation across ages of an ontology in the ApoE4 mice. Lastly, ion-transporters are upregulated in male and female ApoE4 mice **(Figure 6B)**. Of note, while downregulated in young ApoE4 and ApoE2 mice, only aged ApoE4 mice display continued dysregulation of *P-type ion-transporters*. Together, these results suggest a shift toward reduced metabolic and regulatory proteomic signatures, apart from ion-transport.

### Impaired Clearance Contributes to Progressive Mitochondrial Dysfunction

Consistent with literature, aged ApoE4 mice exhibit hallmarks of impaired mitochondria and vesicle processing. As observed in young mice, SNARE proteins are altered in aged ApoE4 mice **(Figure 7A)**. Though young mice experienced a reduction in SNARE proteins, aged mice display a significant increase. Notably, aged female and male mice experience more significant membrane trafficking ontological changes than young mice, though only SNARE complexes are significant for both sexes. Our data show males are significantly altering regulation at the fusion and exocytosis level (with increased synthesis) while females are regulated more upstream at endocytosis and lysosome generation (increased degradation) **(Figure 7C)**. Interestingly, aged female ApoE4 mice reflect a distinct downregulation of V-type ATPases—machinery necessary for vesicle maturation **(Figure 7A).**

**Figure 7.**
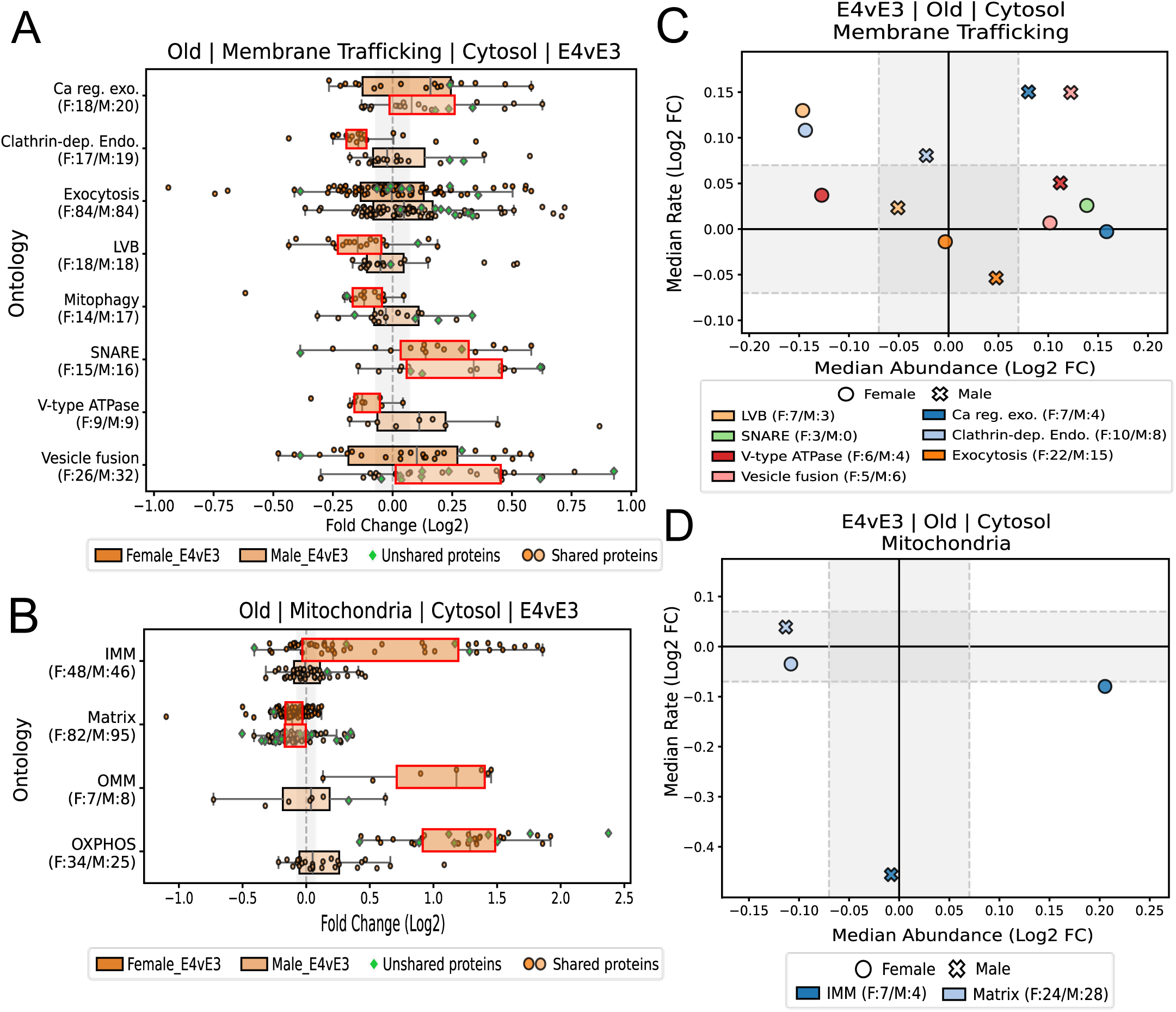
Aged ApoE4 Mice Exhibit Accumulated SNARE Machinery and Mitochondrial Matrix Dysregulation. Boxplots of normalized protein abundances of 18-month-old mice organized into functional groups for female (top, dark) and male (bottom, light) in **(A)** Membrane Trafficking, **(B)** and Mitochondrial Ontologies. Proteostasis Plot of median log2FC abundances (x-axis) and rate (y-axis) for **(C)** Membrane Trafficking and **(D)** Mitochondrial ontologies that exceeded the fold-change cutoff, had p-value significance, and more than 2 proteins with rates. For **(A)** and **(B)**, X-axis for is log2 fold change of ApoE4/ApoE3, y-axis for A and B are ontologies. Fold Change cutoffs for abundance and rate ontologies are |0.07|. Red outlines on boxes represent a significant Wilcoxon p-value of less than 0.05. Ca reg. exo. = Calcium-ion regulated exocytosis, Clathrin-dep. Endo. = Clathrin-dependent endocytosis, LVB = Lysosome Vesicle Biogenesis, Mitophagy = Autophagy of Mitochondrion, IMM = Inner Mitochondrial Membrane, Matrix= Mitochondrial matrix, OMM = Outer Mitochondrial Membrane, OXPHOS= Oxidative Phosphorylation.

Unexpectedly, reduction of *mitochondrial matrix* proteins is the only sex-conserved ontological change in aged E4vE3 mitochondrial ontologies **(Figure 7B)**. This result is surprising as OXPHOS were significantly affected in young mice and play a larger role in cellular energy generation than mitochondrial matrix proteins. In contrast, females additionally express an increase of IMM, OXPHOS, and OMM proteins. This is consistent with the non-uniform membrane to matrix relationship experienced in young ApoE4 female mice. Rate data on the IMM suggests this increase is driven by decreased degradation, indicating retained mitochondrial proteins **(Figure 7D)**. Though most ApoE4 cytosolic ontological abundances have varied dramatically between old and young mice, the membrane fractions of aged mice remain stable **(Supplementary Figure 3-5).**

### ApoE2 Sustains ApoE3-like Protein Homeostasis

To understand how the ApoE2 isoform confers protection against Alzheimer’s Disease, we evaluated the same ontological processes interrogated in ApoE4 mice. Interestingly, ApoE2 exhibits a notable absence of significance for selected ontologies in young **(Figure 8)** mice in the cytosolic fraction, with the exception of *Ion transport by P-type ATPases*. However, this ontology was also significantly reduced in young ApoE4 mice and is therefore not considered a driving factor of isoform-specific change. Remarkably, aged ApoE2 mice also share ApoE3-like expression of proteins in the cytosolic and membrane fraction **(Supplementary Figure 6, 10-12).**

**Figure 8.**
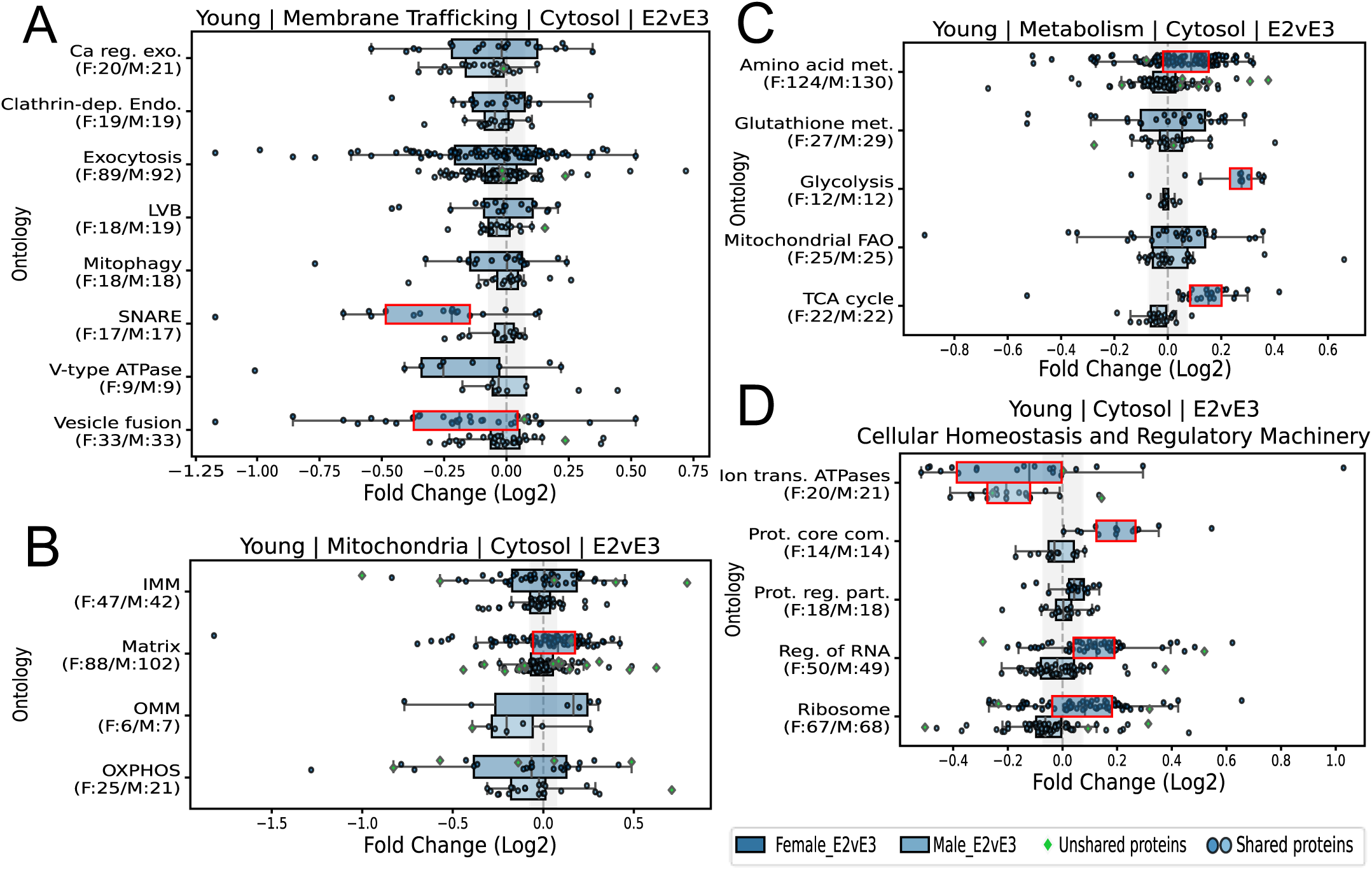
ApoE2 Maintains Proteomic Equivalence to ApoE3 Across Major Functional Ontologies. Boxplots of normalized protein abundances of 3-month-old mice organized into functional groups for female (top, dark) and male (bottom, light) in **(A)** Membrane Trafficking, **(B)** Mitochondria, **(C)** Metabolism, and **(D)** Cellular Homeostasis and Regulatory Machinery Ontologies. X-axis is log2 fold change of ApoE2/ApoE3, y-axis for are ontologies. Fold Change cutoffs for abundance and rate ontologies are |0.07|. Red outlines on boxes represent a significant Wilcoxon p-value of less than 0.05. Ca reg. exo. = Calcium-ion regulated exocytosis, Clathrin-dep. Endo. = Clathrin-dependent endocytosis, LVB = Lysosome Vesicle Biogenesis, Mitophagy = Autophagy of Mitochondrion, IMM = Inner Mitochondrial Membrane, Matrix= Mitochondrial matrix, OMM = Outer Mitochondrial Membrane, OXPHOS= Oxidative Phosphorylation, Amino acid met. = Amino acid metabolic process, Glutathione met. = Glutathione metabolism, Glycolysis = Canonical Glycolysis, Mitochondrial FAO = Mitochondrial fatty acid beta-oxidation, TCA cycle = Tricarboxylic acid cycle, Ion trans. ATPases = Ion transport by P-type ATPases, Prot. Core com. = Proteasome core complex, Prot. Reg. part. = Proteasome regulatory particle, Reg. of RNA = Regulation of RNA splicing.

**Figure 9.**
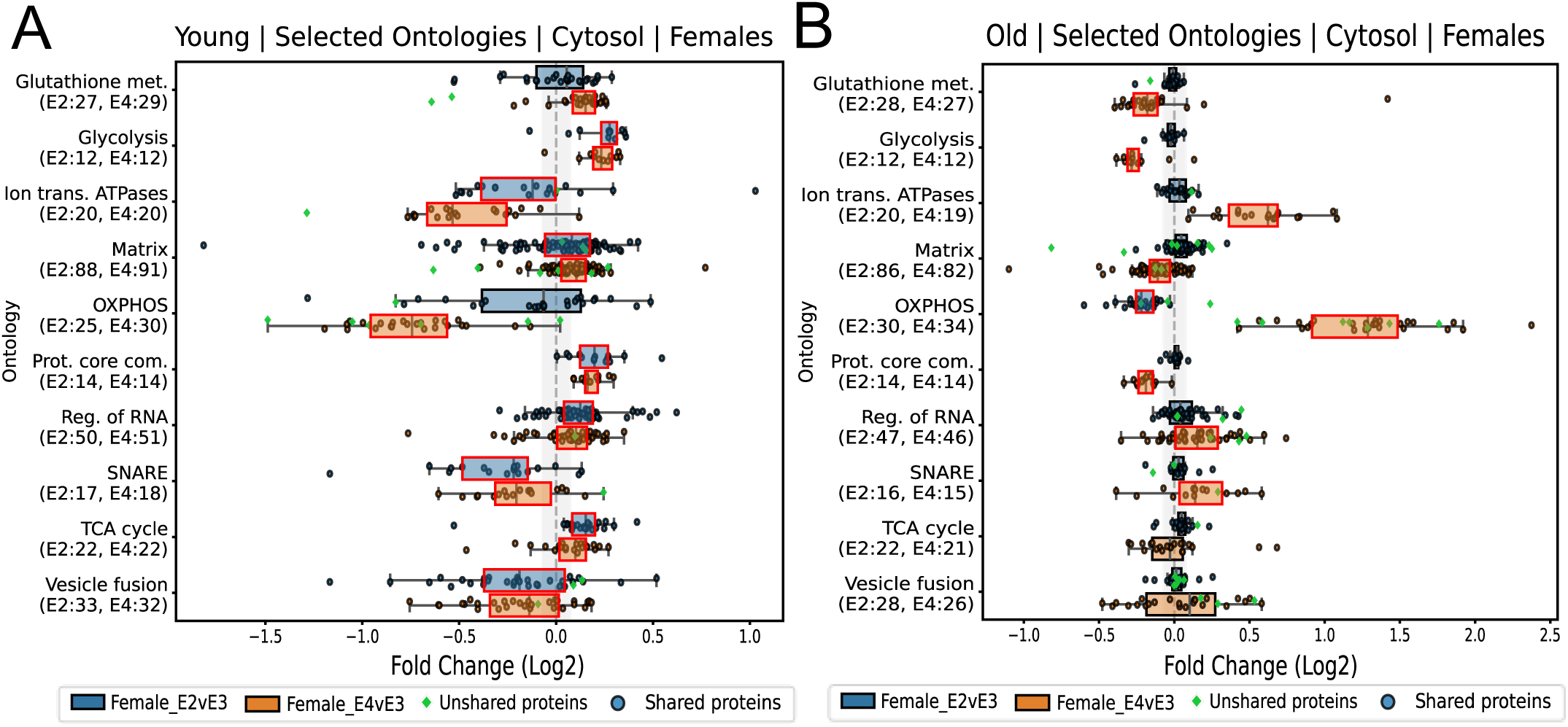
Convergence of Early-Life Proteomic Shifts in Apoe2 and Apoe4 Females Suggest a Sex-Linked Genotype Dependent Response. Boxplots of normalized protein abundances organized into functional groups for **(A)** 3-month-old and **(B)** 18-month-old female mice. This mainly includes ontologies that were equivalently shifted in E2 (top, blue) and E4 (bottom, orange) females, as well as two additional ontologies not changing in E2 that are changed in E4. Displaying the lack of E2 inverse relationship to E4. X-axis is log2 fold change of ApoE2/ApoE3 (Blue) and ApoE4/ApoE3 (Orange), y-axis are ontologies. Fold Change cutoffs for abundance and rate ontologies are |0.07|. Red outlines on boxes represent a significant Wilcoxon p-value of less than 0.05. Glutathione met. = Glutathione Metabolism, Glycolysis = Canonical glycolysis, Ion trans. ATPases = Ion transport by P-type ATPases, Matrix= Mitochondrial matrix, OXPHOS= Oxidative Phosphorylation, Prot. Core com. = Proteasome core complex, Reg. of RNA = Regulation of RNA splicing, TCA cycle = Tricarboxylic acid cycle.

**Figure 10.**
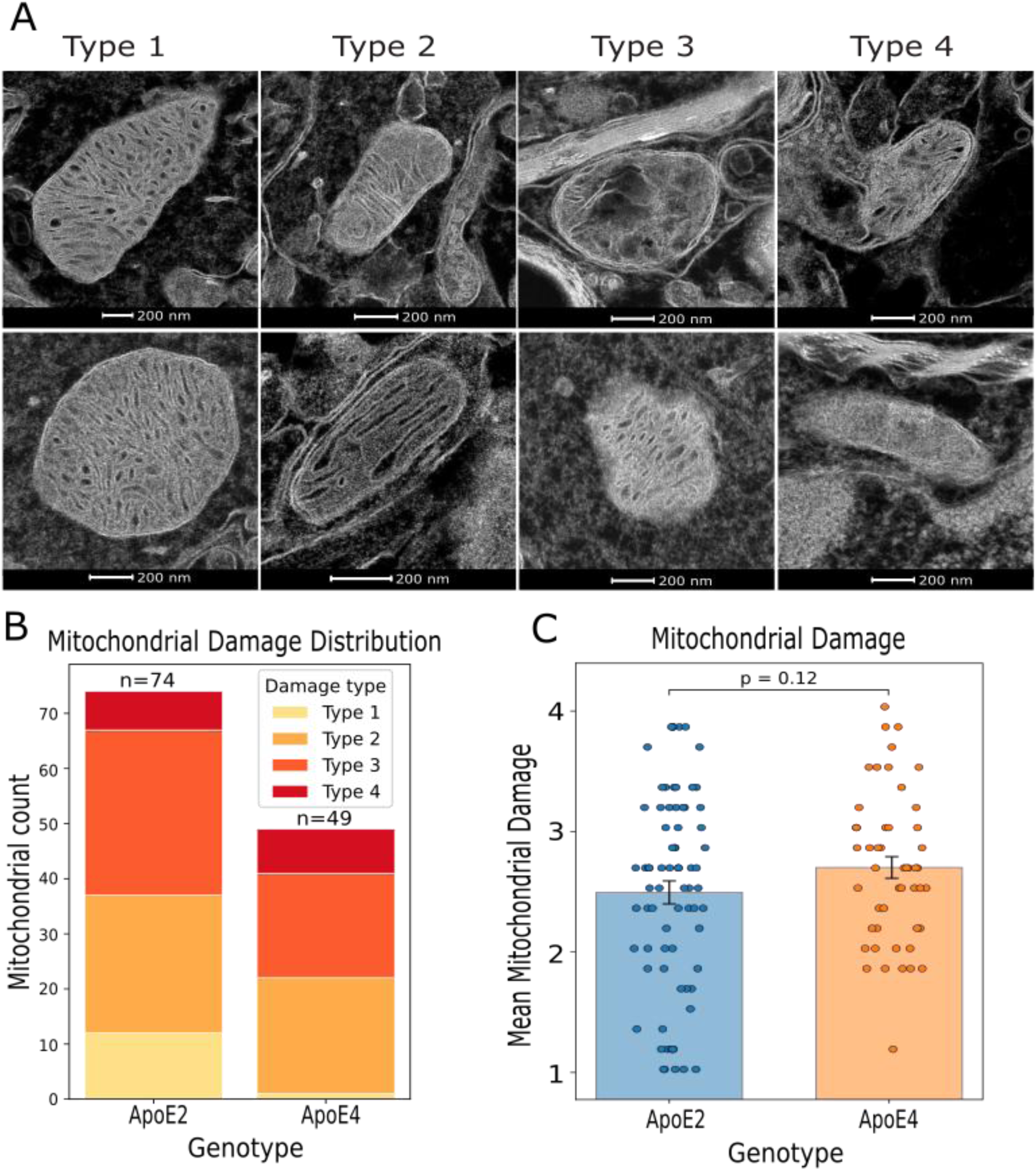
**Mitochondrial Functionality Assessed by Structural Damage Indicates no Genotypic Differences**. **(A)** Mitochondrial damage in old hippocampal tissue was assessed using a scale of 1-4, indicating the severity or type of damage. Results of analysis depicted **(B)** by breakdown of type of mitochondrial damage and **(C)** Mean ± SEM of mitochondrial damage for ApoE2(blue, n=74) and ApoE4(orange, n=49). Error is indicated as SEM of 2 replicates for each genotype, with 74 mitochondria for ApoE2 and 49 mitochondria for ApoE4. Type 1 (healthy/functional): Organized cristae and distinct double membrane, Type 2 (early damage): Loss of dense and organized cristae, membrane still intact. Type 3 (advanced damage): Cristae fragmentation and homogenization and possible discontinuous double membrane. Type 4 (dysfunctional): Major cristae loss and possible mitochondrial rupture.

Together, these data show a lack of ApoE2-specific significant changes in the selected ontologies.

### ApoE2-Mediated Protection is not a Direct Inverse of ApoE4-Driven Changes

While the data in **Figures 4-7** characterize ApoE4-dependent dysregulation and adaptations, **Figure 8** demonstrates a lack of change in ApoE2. This suggests ApoE2-mediated protection is not a direct inverse of ApoE4-driven changes. While there were no detected ApoE2-isoform specific ontological changes, we did observe unique changes within the young female cytosolic **(Figure 8)** and membrane fractions **(Supplementary Figure 7-9)** of ApoE2 mice. ApoE2 and ApoE4 female mice experience similar ontological shifts in early age in the cytosolic fraction **(Figure 9A),** an additional example of the non-linearity between ApoE4 and ApoE2 proteostasis. Notably, these changes are progressively altered in aged ApoE4 mice but stabilize in ApoE2 female mice **(Figure 9B)**.

### Mitochondrial Damage is Unchanged Between Aged Female ApoE2 and ApoE4 Mice

To provide a functional analysis alongside our proteomic observations, we performed a morphological assessment of mitochondrial structural damage; as mitochondrial structure is highly related to its function^44,45^. Based on a 4-tiered scale of mitochondrial damage **(Figure 10A)** inspired by Shults^22^ and Rybka et al.^23^, transmission electron microscopy (TEM) images of aged female brain mitochondria were assessed by a panel of six blinded reviewers. Although ApoE2 mice exhibited a greater proportion of healthy mitochondria (Type I) and ApoE4 mice showed a tendency toward more advanced stages of mitochondrial structural damage, a Welch’s t-test comparing mean mitochondrial damage scores between genotypes revealed no significant difference in overall structural damage **(Figure 10B–C).**Together, this shows a diverse range of damage in a subpopulation of aged mitochondria in both genotypes (ApoE2 and ApoE4).

## Discussion

ApoE4-dependent slowing of endolysosomal processing^14,41,46,47^ and reduced mitochondrial quality^11,15,48–53^ have been well documented in literature. These two processes are highly interconnected, where dysfunction in one leads to dysfunction in the other^13,17^. While ApoE4-induced dysfunction of both pathways is well-supported, neither has been established as the initiating driver of ApoE4-specific proteostasis decay. Our experimental design uses a subtle mouse model that does not experience neurodegeneration to investigate the earliest ApoE and gender-specific proteostasis changes in membrane and cytosolic fractions. In these healthy mice, we find simultaneous early changes in regulation of mitochondrial and vesicle processing, followed by compensatory metabolic rewiring^1^ in young ApoE4 mice **(Figure 11, right)**. For ApoE2 mice, we find striking similarity to ApoE3 mice in young and old cohorts and thus present it as a healthy functioning model **(Figure 11, 12** **(left)).**

**Figure 11.**
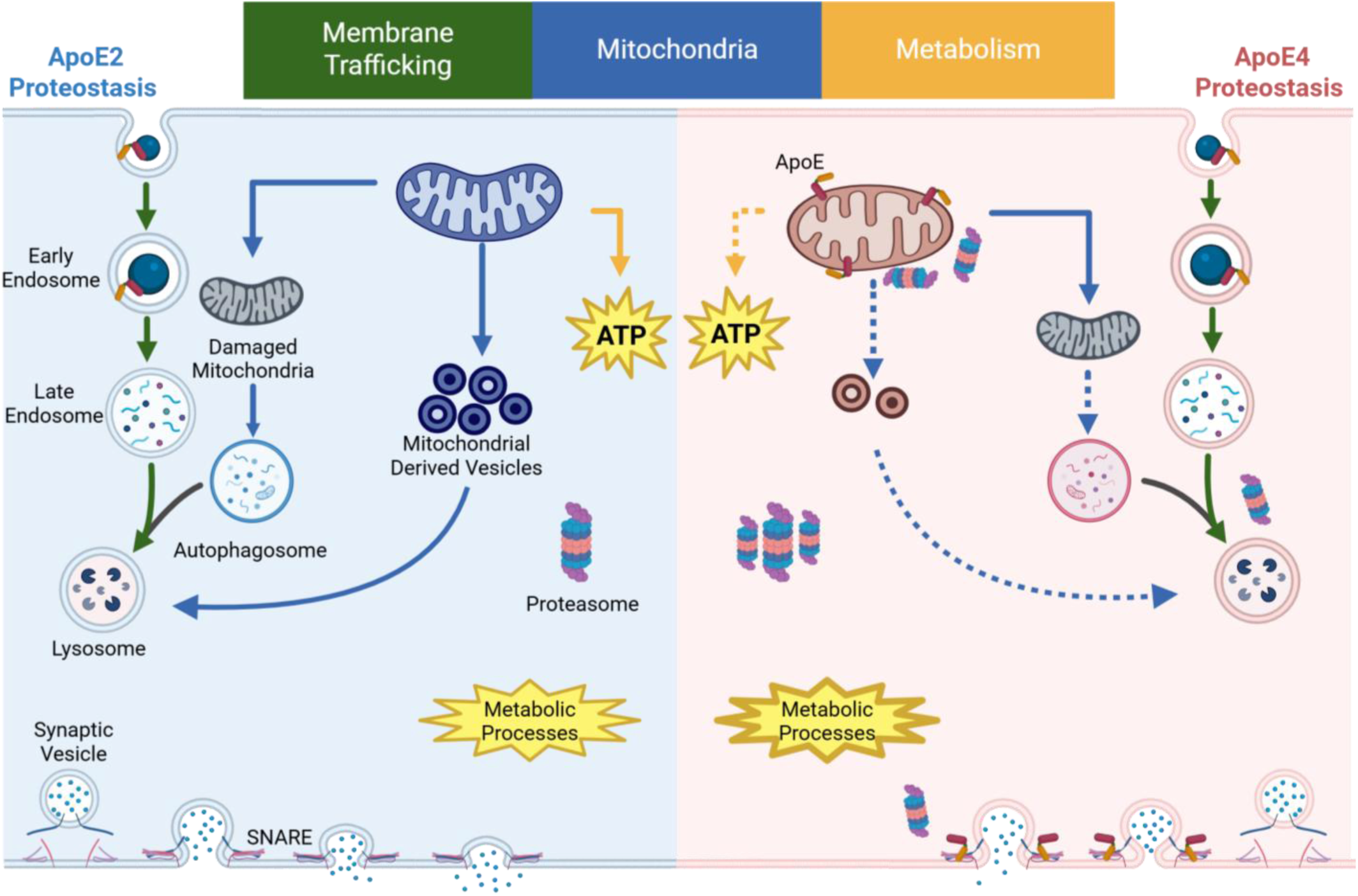
**ApoE4 Young Female Mouse Proposed Mechanism of Hyperactivity**. ApoE2 maintains similarity to healthy ApoE3 controls (left), while ApoE4 experiences significant dysregulation (right). Simultaneous dysfunction of vesicle trafficking and mitochondrial maintenance is initiated by intercalation of ApoE4 into mitochondrial membranes and between SNARE machinery. Dashed arrows and outlines indicate reduction of process. Arrows are coordinating colors with the green, blue, and yellow headers at the top of the image. This proposed mechanism is based off female E4vE3 proteostasis due to its apparent increased AD pathological progression compared to males.

**Figure 12.**
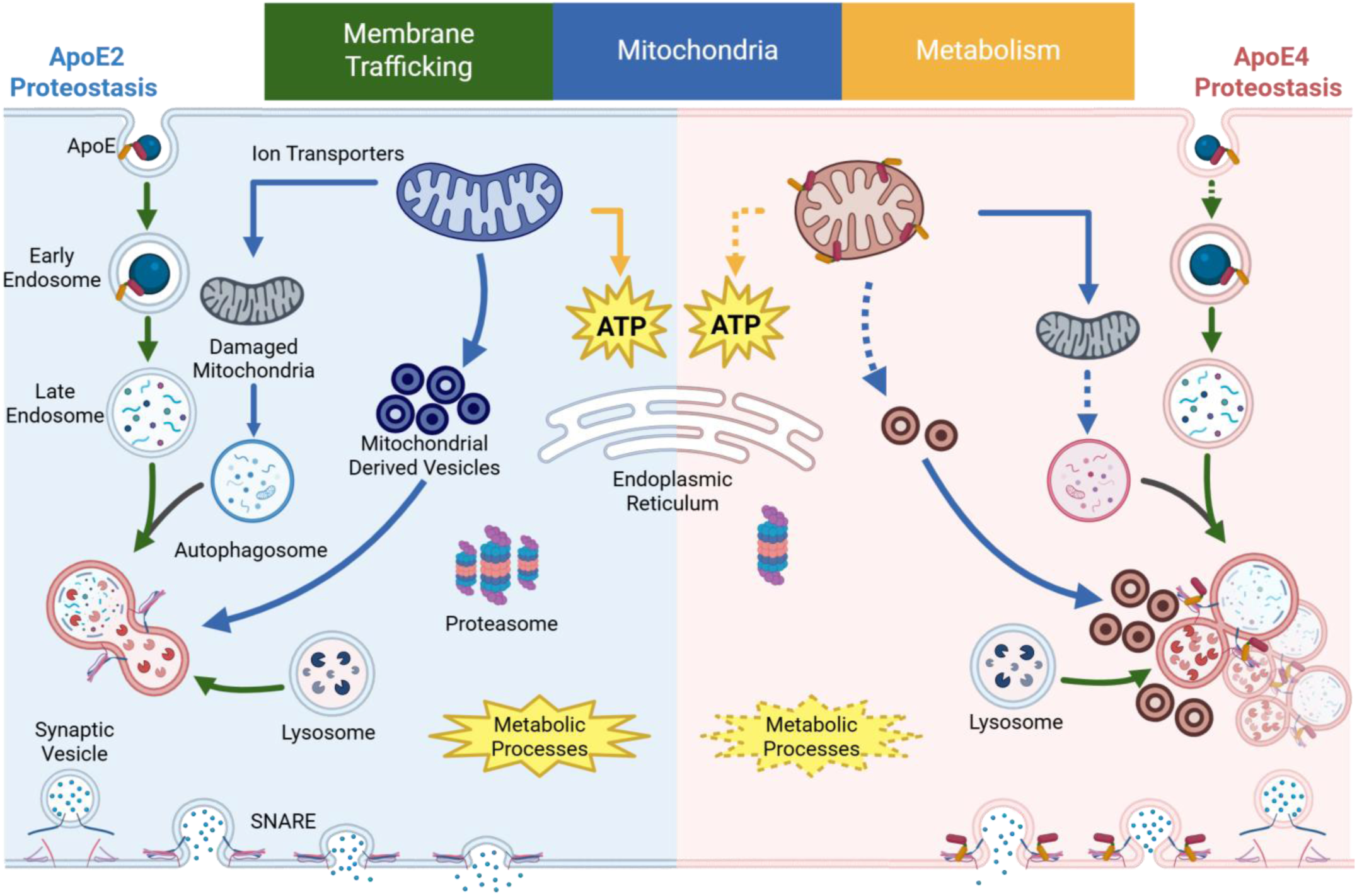
**ApoE4 Aged Female Mouse Proposed Mechanism of Exhaustion**. Dashed arrows and outlines indicate reduction of process. Arrows are coordinating colors with the green, blue, and yellow headers at the top of the image. This proposed mechanism is based off female E4vE3 proteostasis due to its apparent increased AD pathological progression compared to males.

Though ApoE2 mice overall experience little change compared to ApoE3 in the selected ontologies, young ApoE2 and ApoE4 female mice exhibit similar ontological shifts **(Figure 9A)**, as observed by Zuniga et al.^3^. Many of these shifts were additionally altered in young ApoE4 males, and aged ApoE4 mice, contributing to our reasoning for discussing them as major ApoE4-dependent ontological changes but not for ApoE2 (see *Data Interpretation Model* in Results). However, the similarities in young female mice reveal a more complex AD risk assessment than protein abundance changes alone can predict. It is well known that ApoE isoforms carry distinct lipid cargo^54,55^, which can result in differing metabolic dependencies and membrane fluidity.

Therefore, despite exhibiting similar proteomic changes, ApoE2 and ApoE4 mice may experience different biological consequences. For example, SNARE proteins are reduced in both genotypes in young females. However, ApoE4 peptides have been shown to impare the functionality of SNAREs^56,57^. Therefore, though reduced SNARE abundance may not be a cause of dysfunction, reduced SNAREs in combination with interrupted functionality could result in more extreme consequences in ApoE4 mice. Our data support the existence of distinct biological environments between ApoE2 and ApoE4, evidenced by opposing kinetic regulation for ion transport, vesicle fusion, RNA regulation, and glycolysis proteins **(Figure 4, 5, and Supplemental Figure 7).** Collectively, these findings suggest that young females are particularly responsive to ApoE-specific environments, and that similar proteomic alterations can produce distinct biological consequences depending on the surrounding metabolic and regulatory context.

In young ApoE4 cytosolic fractions, our findings support a model in which the physical intercalation of ApoE4 disrupts vesicle processing^41^ and mitochondrial quality^58^ simultaneously. The reduction of *calcium-ion regulated exocytosis* and *SNARE proteins* in combination with ApoE4 peptide interference potentially impairs the ability of the cells to adequately fuse vesicles^41,58–62^ and transmit signals as frequently observed in ApoE4 mouse models **(Figure 4A)**^57^. While related studies that identified ApoE4 interactions with SNARE machinery found SNARE abundance unchanged^41^, our model suggests the disrupted fusion occurs in combination with a decrease of SNARE proteins specifically residing in the cytosolic fraction of young mice. Simultaneously, the ApoE4-specific decline in oxidative phosphorylation (OXPHOS) complexes^63^ may be a response induced by ApoE4 peptide insertion into the mitochondrial membrane which has previously been observed^1,10,49,50,56,58,64,65^ **(Figure 4B)**. We propose this mechanical disruption not only begins to create physical blockage of protein transport, but also destabilizes the mitochondrial membrane potential, contributing to the mitochondrial matrix depletion observed in aged mice^13^ **(Figure 7B)**. Taken together, these early disruptions support a simultaneous (as opposed to sequential) dysregulation of vesicle and mitochondrial processing.

Though males and females both report these early ApoE4-peptide dependent decreases in SNARE machinery, their kinetic mechanisms of membrane trafficking ontologies differ. ApoE4 males display increased degradation, while females report decreased synthesis **(Figure 4C)**. In cellular regulation and homeostasis ontologies, similar disconnects in sex-specific regulation are observed in young mice **(Figure 5C)**. This indicates that ApoE4 related dysfunction may be due to changes in protein abundance, not the mechanism used to achieve it. This also suggests that though pathological dysfunction due to ApoE4 has similar characteristics in males and females, treatment should be specifically tailored for sex.

Although a reduction in OXPHOS^1,6,13,50,51,54,62^ has been observed numerous times in literature reflecting the ApoE4 induced metabolic stress, our female data suggests the reduction of OXPHOS may not solely be a consequence of metabolic insult. Because OXPHOS complexes primarily reside in the IMM, their reduction alongside broader reductions in IMM and OMM proteins in young females suggests an overall mitochondrial membrane perturbation. Interestingly, this is not accompanied by a reduction of matrix proteins, suggesting a physical restructuring of the mitochondria versus whole mitochondrion clearance **(Figure 4B)**. Notably, the membrane fraction is not significantly changed in any of these three mitochondrial components compared to ApoE3 **(Supplemental Figure 1),** suggesting the patterns observed in the cytosolic fraction cannot be explained by protein retention in the membrane and thus are indeed a phenomenon localized to the cytosol. A similar disconnect between changes in mitochondrial membrane versus changes in mitochondrial matrix was observed previously in transgenic ApoE4 mice^3^.

The cytosolic rearranging of the mitochondria could be related to mitochondrial-derived vesicles (MDVs in **Figure 11**) ^17,66,67^. We propose the mitochondria in ApoE4 mice attempt to prune away damaged complexes via MDVs^17^ consisting of inner (IMM) and outer mitochondrial membrane (OMM) components^68^, but are unable to do so at the same pace as healthy ApoE3 mice^66,69,70^. The reduction of MDVs due to impaired pruning results in a relative decrease of cytosolic OMM and IMM components^14,68,70^, as well as the accumulation of unhealthy mitochondria that swell with matrix components^1^ **(Figure 4B)**. In male mice, we propose the decline in MDV-mediated mitochondrial pruning is delayed, with the current dataset capturing the advanced pathological progression in females^40,51^. Alternatively, the sexual dimorphism observed in regulation may be extrapolated to suggest that while reduced MDV budding is the female response to ApoE4 induced damage, males compensate for the mitochondrial membrane dysfunction by other means and thus never achieve the level of damage observed in females.

The early proteostatic disruptions of vesicle trafficking and mitochondrion in young ApoE4 mice induce significant metabolic adaptation. With the reduction of OXPHOS complexes and concomitant increase in glycolytic enzymes due to reduced degradation **(Figure 5A, C)**, metabolism likely shifts toward increased aerobic glycolysis^13,71^ to maintain energy demands. Similar shifts in metabolism have been observed in healthy young human brains^49,69,72^. Though still healthy, the increased synthesis of glutathione metabolic proteins in young ApoE4 mice indicates an adaptive response toward elevated ROS generation **(Figure 5A,C)**. This is consistent with increased oxidative stress frequently observed in ApoE4 models^5,40,53,73^. Young ApoE4 female mice additionally experience an increase in proteasomal core subunits **(Figure 5B)**, likely to mediate protein clearance in response to interrupted mitochondrial pruning and vesicle processing.

Consistent with the literature, the proteomics suggest an early-life ApoE4-driven hyperactivity that results in late-stage exhaustion^52,74–77^. We observed a proteomic signature of reduced metabolic effort in aged ApoE4 mice **(Figure 12, right)**. The compensatory elevation of glycolysis and amino acid metabolism^51^ observed in young mice is replaced by a systemic decline in protein abundance for these ontologies^62,78^, which is further compounded by a reduction in fatty acid beta oxidation **(Figure 6A)**. Notably, aged female ApoE4 mice additionally exhibit a reduction in glutathione metabolism further supporting the enhanced AD pathological progression for females expressing ApoE4^79^. While previous studies have also reported a sex-specific altered glutathione metabolism^80,81^, our study reveals significantly altered glutathione metabolism in old ApoE4 females instead of males^40^ . We suspect this is due to differences in the type of measurement or our ApoE isoform model in contrast to AD or healthy gender-specific studies. Females additionally exhibit an interesting regulation of the proteasome, with reduced synthesis for proteasome regulatory particles and increased degradation of the proteasome core, displaying a lack of regulatory coordination within the complex **(Figure 6D)**. Reduction of proteasome complexes and loss of complex stoichiometry has been associated with aging, cellular stress, and reduced activity^82–85^. Collectively, these data suggest reduced metabolic effort and loss of regulatory coordination in aged ApoE4 mice.

In addition to reduced metabolic ontologies, we observe continued differential mitochondrial proteome signatures in aged ApoE4 mice compared to ApoE3. Notably, the conserved mitochondrial signature in both ApoE4 males and females is a cytosolic attenuation of the matrix proteome **(Figure 7B)**. This decline encompasses a significantly broader range of proteins than the specific TCA cycle reductions reported in previous studies^13,51,71^, suggesting a more comprehensive loss of mitochondrial content. We investigated the decreased matrix components to determine whether a specific functional category could explain the phenotype. However, we were unable to define any apparent relationship between the decreased proteins and their size, signal sequence, function, or translation^86^ **(Supplementary Figure 13)**. This argues against mitochondrial leakage, selective import, biological utility, or genomic origin as the primary drivers of the observed matrix discrepancy. This leads us to suspect we are seeing additional support for ApoE4 peptides disrupting membrane complexes necessary for import of matrix proteins^17^ and stabilization of mitochondria. The peptides may also be disrupting the membrane potential^62^ or ATP availability, ultimately impairing mtDNA translation^87–89^. Alternatively, the unique lipid environment established by ApoE4 may be similarly compromising protein import, export, or assembly of protein complexes^62^.

Similar to young ApoE4 mice, aged ApoE4 females additionally display significantly remodeled IMM and OMM proteins in the cytosolic fraction **Figure 7B)**, relating to an MDV signature. However, in young mice there was an apparent reduction in MDVs whereas in old mice the MDVs have accumulated^47^ in the cytosol. We suspect this accumulation is due to an age-dependent progressive impairment of SNARE-mediated vesicle clearance^90^, leading to a steady accrual of MDVs over time. This is supported by an observed decrease in the degradation of IMM components **(Figure 7D).**

To investigate whether the proteomic signatures of ApoE4-mediated dysregulation in the cytosol are impacting the mitochondrial structure and function^91^, we evaluated mitochondrion structural quality (**Figure 10**). We used a 4-tiered scale of structural damage focusing on cristae density and membrane integrity, as mitochondrial functionality is closely linked to cristae organization^44,45,92,93^ and the double membrane^94^. Our analysis in the hippocampus of aged female mice demonstrate that mitochondria in ApoE4 mice trend toward advanced stages of damage, though the aggregate mean score for mitochondrial damage did not reach statistical significance (p = 0.12) when compared to the ApoE2 genotype. Furthermore, the distribution of mitochondrial damage between ApoE2 and ApoE4 is similar, indicating our proteomic analysis is measuring dysregulation that cannot be assessed via whole mitochondria morphological changes. While not significant, a bias within assembled mitochondria (**Figure 9**), in aged ApoE4 mice may indicate a trend toward advanced aging in which the mitochondrial population is increasingly damaged^95^. Ultimately, these results indicate that the proteomic ApoE4-mediated pathology is not expressed as whole mitochondrial structural damage and overall dysfunction, further supporting changes in small-scale quality control such as MDV release.

To fully assess isoform-specific proteostasis changes, we extended our analysis to include ApoE2 cohorts. The only significant change shared by male and female young ApoE2 mice was a reduction in ion transport by P-type ATPases **(Figure 8)**, a change convergent with our observations in young ApoE4 mice (**Figure 5B**). Of note, while ApoE2 proteostasis for ion transport in old mice is no longer significantly changed, ApoE4 mice continue to experience dysregulation. Therefore, we suggest ion transport related changes are not a driving factor of young ApoE4 dysregulation, but are an important characteristic of aged ApoE4 mice^12^. Importantly, these data suggest that the significant proteostatic changes observed in ApoE4 mice are not mirrored in ApoE2 (non-reciprocity). This indicates that the protective effect of ApoE2 is not simply a reversal of the ApoE4-induced dysfunction and highlights the non-linearity of the relationship among ApoE2, ApoE3, and ApoE4.

Another example of the non-reciprocity is our observation of many similar ontological shifts for young ApoE2 and ApoE4 females relative to ApoE3. Interestingly, while aged ApoE4 mice continually express significance in these ontologies, suggesting a progressive state of dysfunction, ApoE2 mice appear to resolve these proteostasis changes and return to a similar homeostatic state to that of aged ApoE3 mice. Additionally, while few ontologies had significant abundance changes in the membrane fraction of ApoE4, female ApoE2 membranes contain multiple. *Ion transport by P-type ATPases* and *proteasome core complex* ontologies are changed in the cytosolic and membrane fractions in the same direction, indicating a more complete cellular response versus a fraction-dependent one as observed in ApoE4. It has been proposed that ApoE2 mice provide AD protection due to more productive regulation^3^. Though our overall rate analysis suggests all genotypes are essentially equivalent in turnover rate, most of the changed ontologies have faster turnover rates in young ApoE2 mice than ApoE3 **(Supplementary Figure 7 and 9)**. This, along with the other ontologies shifted in ApoE2 female membranes, may suggest the ApoE2 protective effect could potentially be due to enhanced regulation of organelle membranes environments.

## Limitations in the Study

Although our proteomic analysis provides insight into ApoE isoform-associated proteostatic fluctuations, these data do not directly establish whole mechanistic functional impairment. The observed signatures are evidence of altered proteomic regulation rather than absolute cellular dysfunction. Direct measurements of mitochondrial respiration, ATP production or consumption, lysosomal activity, and autophagic flux, would strengthen the mechanistic interpretation of these findings. Additionally, post-translational modifications (PTM), which frequently regulate protein localization and function, were not assessed in this study.

Furthermore, our kinetic analysis was more limited than the abundance-based dataset, reducing the number of proteins and ontologies for which turnover rates could be confidently reported. Finally, this study analyzed whole-brain proteomic fractions and therefore did not resolve cell type-specific regulation, which should be addressed in future cell type-resolved proteomic studies.

## Conclusion

In this study we integrated protein abundance and kinetics to elucidate the mechanisms underlying ApoE isoform-specific proteostasis. Our findings establish critical links between previously isolated observations of vesicle trafficking, mitochondria dysfunction, metabolic changes, proteasome activity, and ion regulation, providing a more complete understanding of ApoE4-driven dysregulation. We propose that early-stage proteostatsis changes are induced by simultaneous compromise of mitochondrial and vesicle processing and is accompanied by a compensatory metabolic hyperactivity. In contrast, the late-stage ApoE4 response is characterized by reduced metabolic and proteasomal effort, disproportionate mitochondria, and accumulation of SNAREs. Furthermore, our data support a non-linear relationship between ApoE2, ApoE3, and ApoE4, offering new insights behind increased risk or protection against Alzheimer’s disease. Finally, we provide evidence of sex-dependent kinetic regulation which modifies the risk of ApoE4-induced damage as well as implying clinical interventions for Alzheimer’s disease may require sex-specific therapeutic strategies. Importantly, our findings demonstrate that ApoE4 independently initiates proteostatic shifts that mirror the hallmark signatures of AD. This suggests that the profound risk associated with the ApoE4 allele stems from a bias in proteome maintenance that enhances the sensitivity to life-style based risk factors even in the absence of amyloid and tau buildup.

## Acknowledgements

We are grateful for the assistance of the BYU Live Animal Facility, the BYU Electron Microscopy facility, and the Fritz B. Burns Biological Mass Spectrometry Facility at BYU. This work was made possible by a grant from the National Institutes of Health [R01AG066874] to JCP; Brigham Young University Undergraduate Research Awards to NP, JME, NEE, BSJ, EGS, MS, RSB, NGM, WD, JGW, ESV, CTG, JRB, JDM, MFP. The funders had no role in study design, data collection and analysis, decision to publish, or preparation of the manuscript

## Data Availability

The mass spectrometry proteomics data have been deposited to the ProteomeXchange Consortium via the PRIDE^96^ partner repository with the dataset identifier PXD079261. This includes the raw LC-MS files used for quantitative and kinetic analysis in this study, as well as the Fragpipe workflow and DIANN outputs.

## Project Name

Ontological Analysis of Brain Proteostasis Highlights the Sex-Dependent Trajectory of ApoE Isoform-Specific Regulation

**Project accession:** PXD079261

**Project DOI:** Not applicable

## Reviewer access details

Log in to the PRIDE website using the following details:

**Project accession:** PXD079261

**Token:** l3a0jbuQvqw1

The code used for data processing is available at https://github.com/aDenos24/ApoE_KI_Proteomics_Denos, as well as the kinetic software (DeuteRater) https://github.com/JC-Price/DeuteRater/releases/tag/DeuteRater-v6-beta

**Supporting Information:** Additional boxplots and proteostasis plots for E2vE3 and E4vE3 (DOC), raw and processed protein abundance and kinetic data (XLSX)

