## Supplemental Figures for "Ontological Analysis of Brain Proteostasis Highlights the Sex-Dependent Trajectory of ApoE Isoform-Specific Regulation"

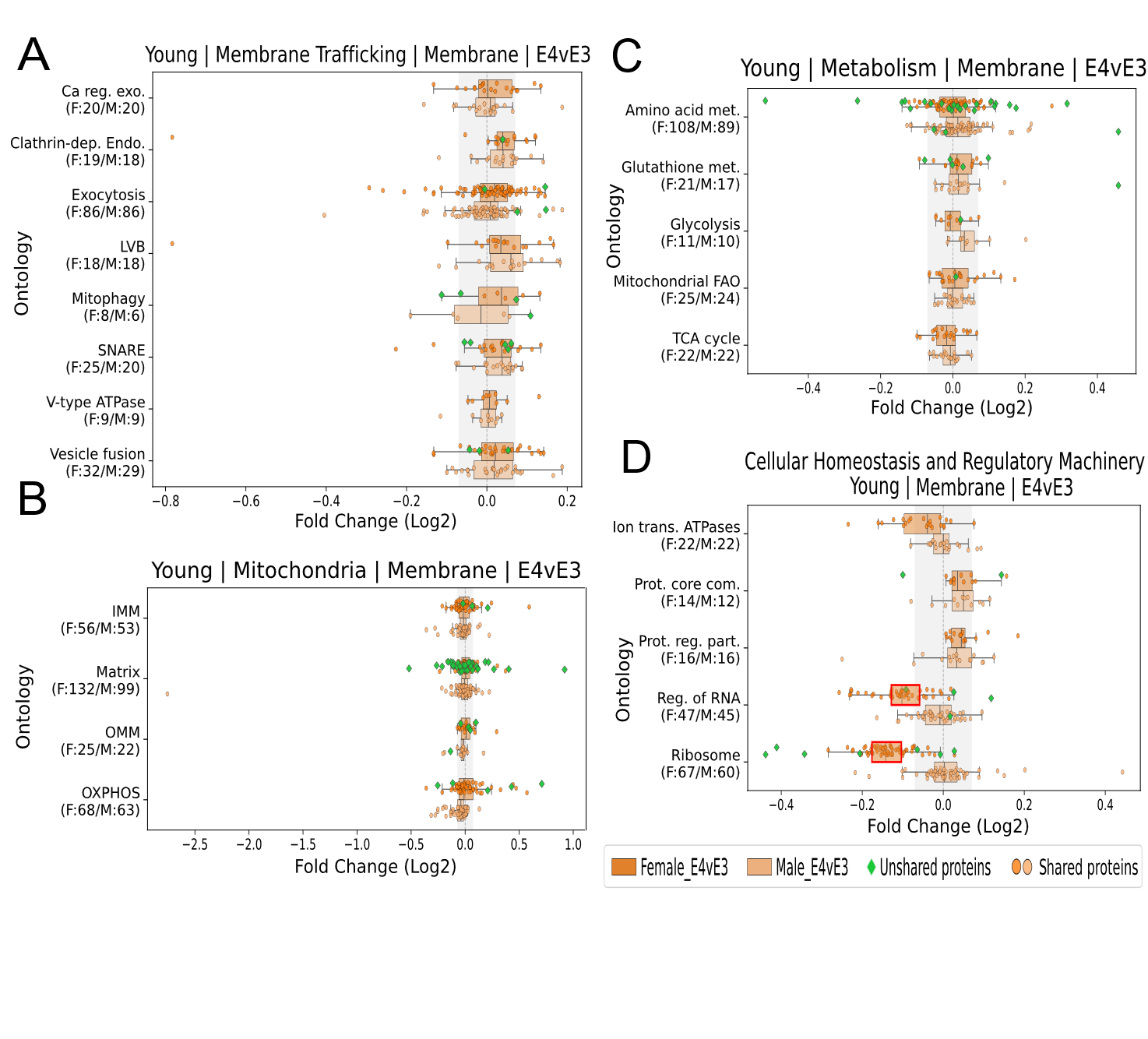

**Supplementary Figure 1. Young E4vE3 Membrane Boxplots**. Boxplots of normalized protein abundances of 3-month-old male and female mice organized into functional groups for female (top, dark) and male (bottom, light) in **(A)** Membrane Trafficking, **(B)** Mitochondria, **(C)** Metabolism, **(D)** and Cellular Homeostasis and Regulatory Machinery ontologies. X-axis is log2 fold change of ApoE4/ApoE3, y-axis are ontologies. Fold Change cutoffs for abundance and rate ontologies are |0.07|. Red outlines on boxes represent a significant Wilcoxon p-value of less than 0.05. Ca reg. exo. = Calcium-ion regulated exocytosis, Clathrin-dep. Endo. = Clathrin-dependent endocytosis, LVB = Lysosome Vesicle Biogenesis, Mitophagy = Autophagy of Mitochondrion, IMM = Inner Mitochondrial Membrane, Matrix= Mitochondrial matrix, OMM = Outer Mitochondrial Membrane, OXPHOS= Oxidative Phosphorylation, Amino acid met. = Amino acid metabolic process, Glutathione met. = Glutathione metabolism, Glycolysis = Canonical Glycolysis, Mitochondrial FAO = Mitochondrial fatty acid beta-oxidation, TCA cycle = Tricarboxylic acid cycle, Ion trans. ATPases = Ion transport by P-type ATPases, Prot. Core com. = Proteasome core complex, Prot. Reg. part. = Proteasome regulatory particle, Reg. of RNA = Regulation of RNA splicing.

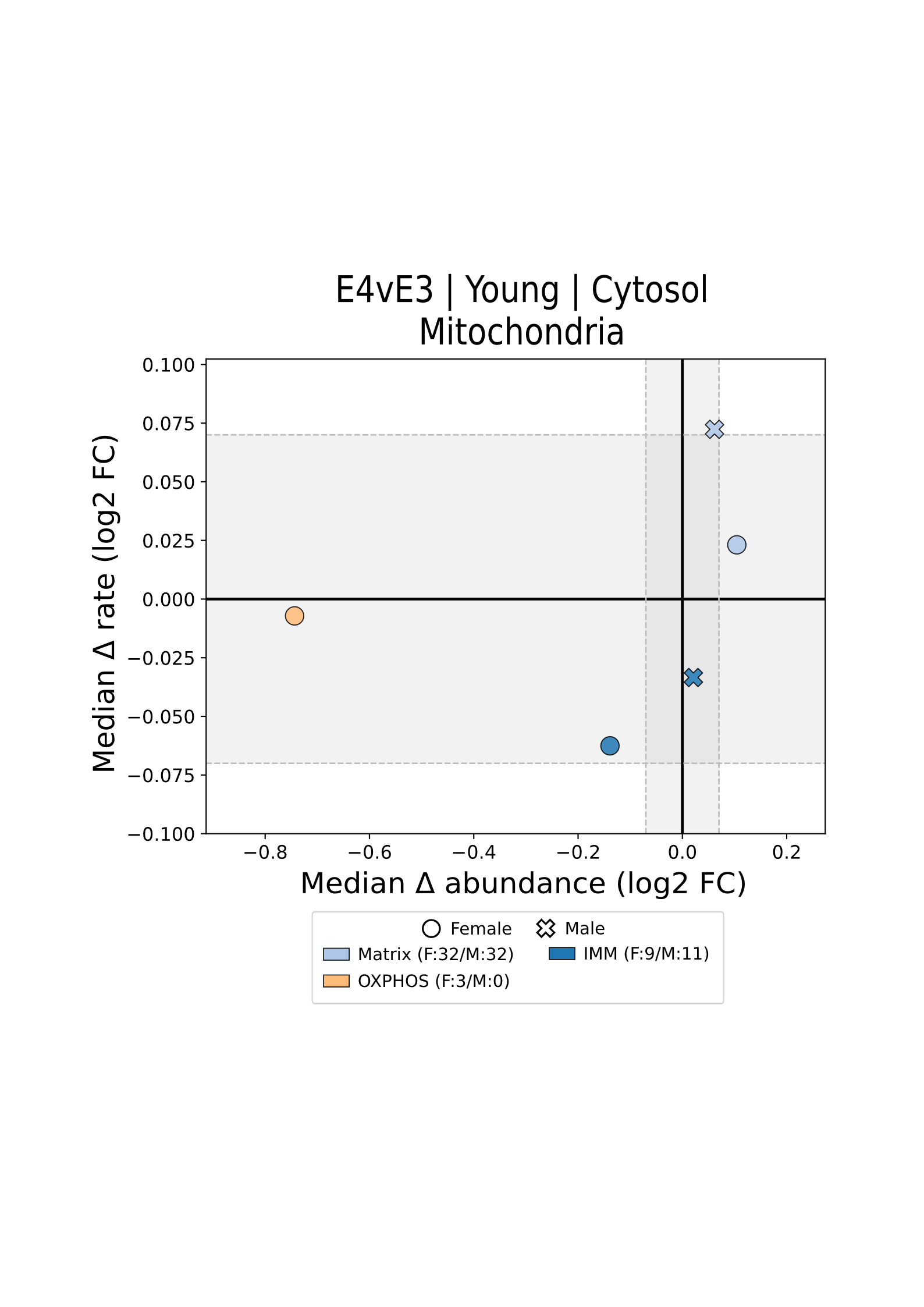

**Supplementary Figure 2. Young E4vE3 Cytosolic Mitochondrial Proteostasis Plot**. Proteostasis plot of normalized protein abundance (x-axis) and rate (y-axis) fold changes of 3-month-old male and female mice organized into functional groups for Mitochondria ontologies. OXPHOS = Oxidation phosphorylation, IMM = Inner Mitochondrial Membrane, Matrix = Mitochondrial Matrix.

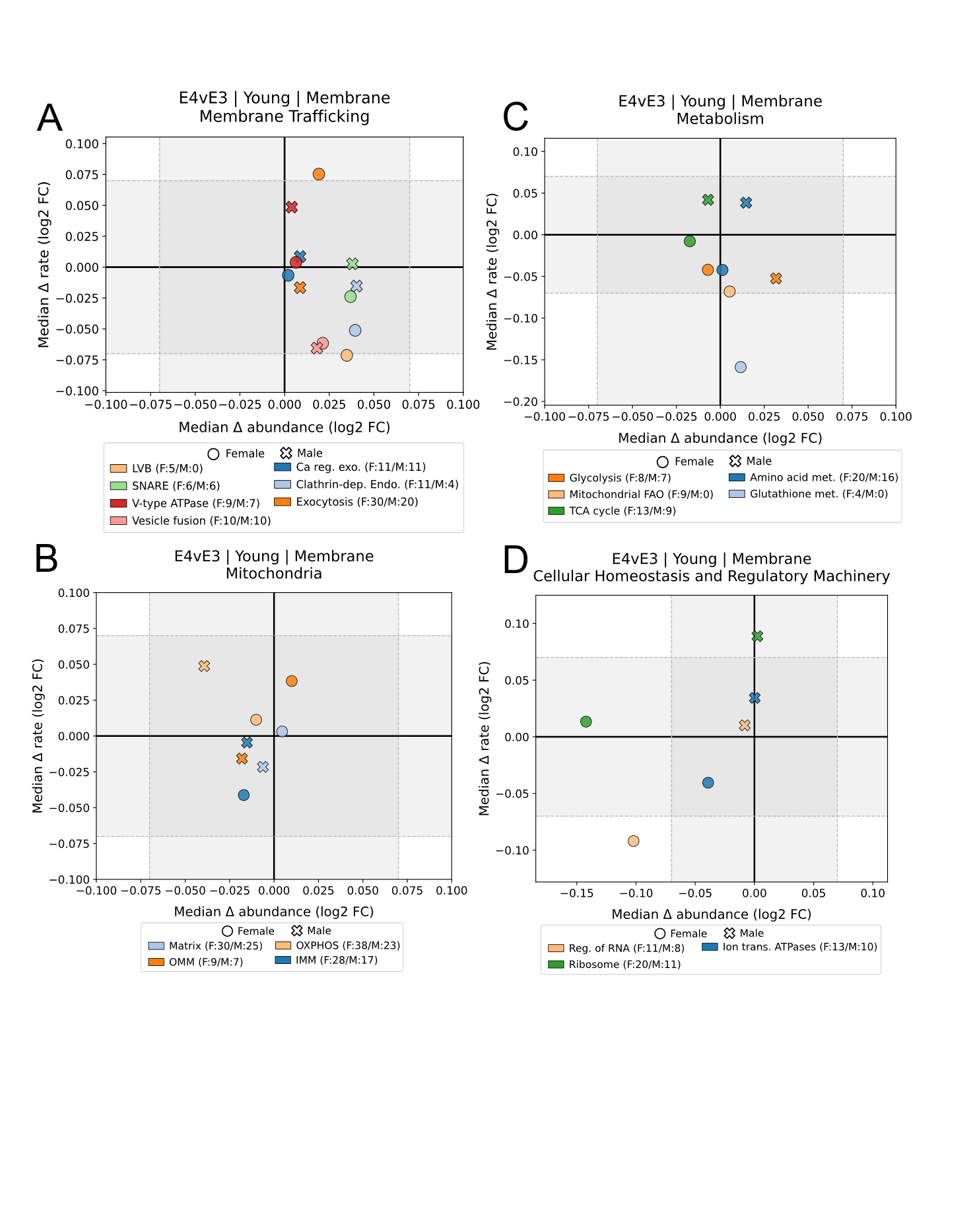

**Supplementary Figure 3. Young E4vE3 Membrane Proteostasis Plots.** Proteostasis plot of normalized protein abundance (x-axis) and rate (y-axis) fold changes of 3-month-old male and female mice organized into functional groups for **(A)** Membrane Trafficking, **(B)** Mitochondria, **(C)** Metabolism, **(D)** and Cellular Homeostasis and Regulatory Machinery ontologies. Fold Change cutoffs for abundance and rate ontologies are |0.07|. Red outlines on boxes represent a significant Wilcoxon p-value of less than 0.05. Ca reg. exo. = Calcium-ion regulated exocytosis, Clathrin-dep. Endo. = Clathrin-dependent endocytosis, LVB = Lysosome Vesicle Biogenesis, Mitophagy = Autophagy of Mitochondrion, IMM = Inner Mitochondrial Membrane, Matrix= Mitochondrial matrix, OMM = Outer Mitochondrial Membrane, OXPHOS= Oxidative Phosphorylation, Amino acid met. = Amino acid metabolic process, Glutathione met. = Glutathione metabolism, Glycolysis = Canonical Glycolysis, Mitochondrial FAO = Mitochondrial fatty acid beta-oxidation, TCA cycle = Tricarboxylic acid cycle, Ion trans. ATPases = Ion transport by P-type ATPases, Prot. Core com. = Proteasome core complex, Prot. Reg. part. = Proteasome regulatory particle, Reg. of RNA = Regulation of RNA splicing.

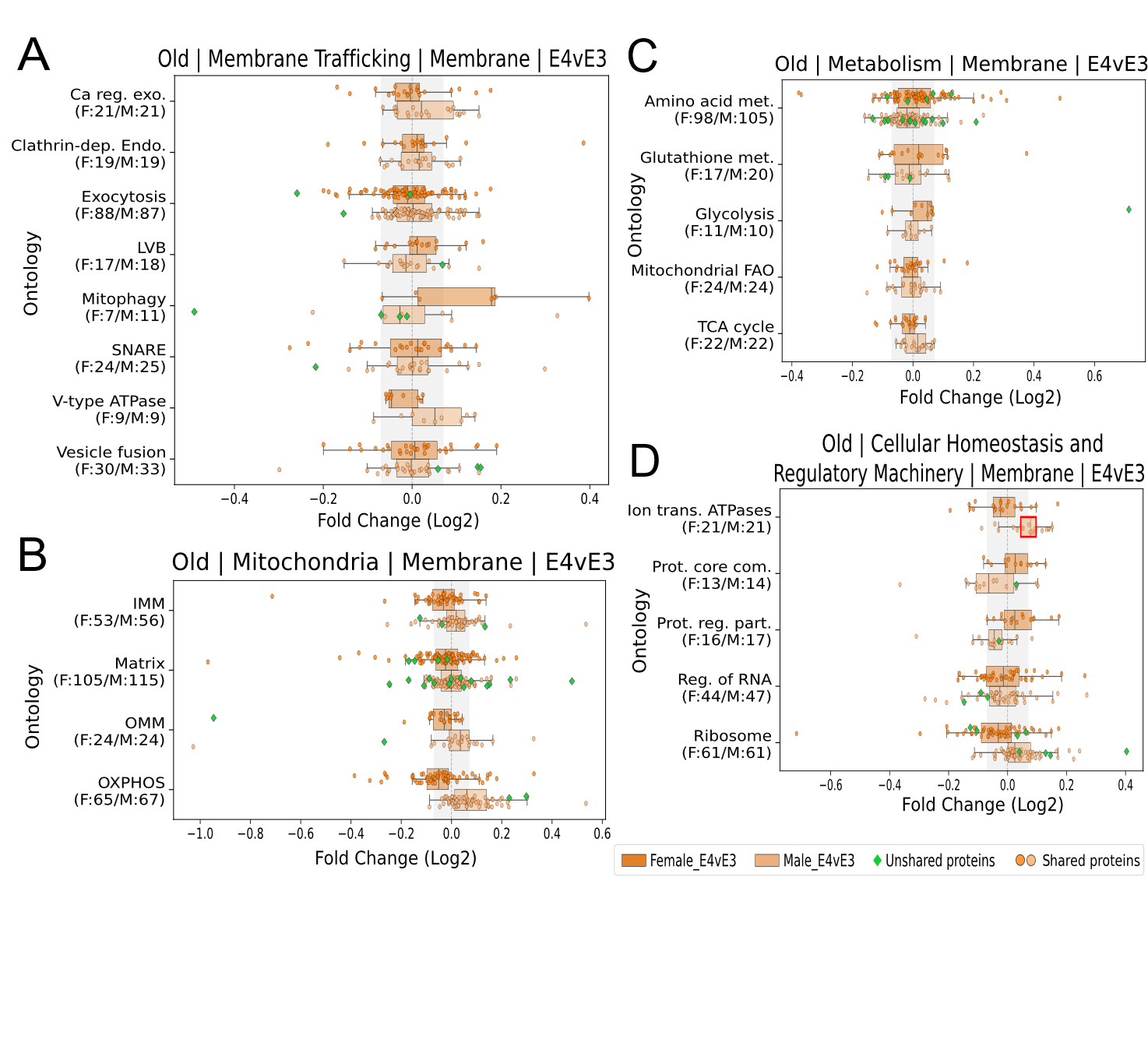

**Supplementary Figure 4. Old E4vE3 Membrane Boxplots.**. Boxplots of normalized protein abundances of 18-month-old mice organized into functional groups for female (top, dark) and male (bottom, light) in **(A)** Membrane Trafficking, **(B)** Mitochondria, **(C)** Metabolism, and **(D)** Cellular Homeostasis and Regulatory Machinery Ontologies. X-axis is log2 fold change of ApoE4/ApoE3, y-axis for are ontologies. Fold Change cutoffs for abundance and rate ontologies are |0.07|. Red outlines on boxes represent a significant Wilcoxon p-value of less than 0.05. Ca reg. exo. = Calcium-ion regulated exocytosis, Clathrin-dep. Endo. = Clathrin-dependent endocytosis, LVB = Lysosome Vesicle Biogenesis, Mitophagy = Autophagy of Mitochondrion, IMM = Inner Mitochondrial Membrane, Matrix= Mitochondrial matrix, OMM = Outer Mitochondrial Membrane, OXPHOS= Oxidative Phosphorylation, Amino acid met. = Amino acid metabolic process, Glutathione met. = Glutathione metabolism, Glycolysis = Canonical Glycolysis, Mitochondrial FAO = Mitochondrial fatty acid beta-oxidation, TCA cycle = Tricarboxylic acid cycle, Ion trans. ATPases = Ion transport by P-type ATPases, Prot. Core com. = Proteasome core complex, Prot. Reg. part. = Proteasome regulatory particle, Reg. of RNA = Regulation of RNA splicing.

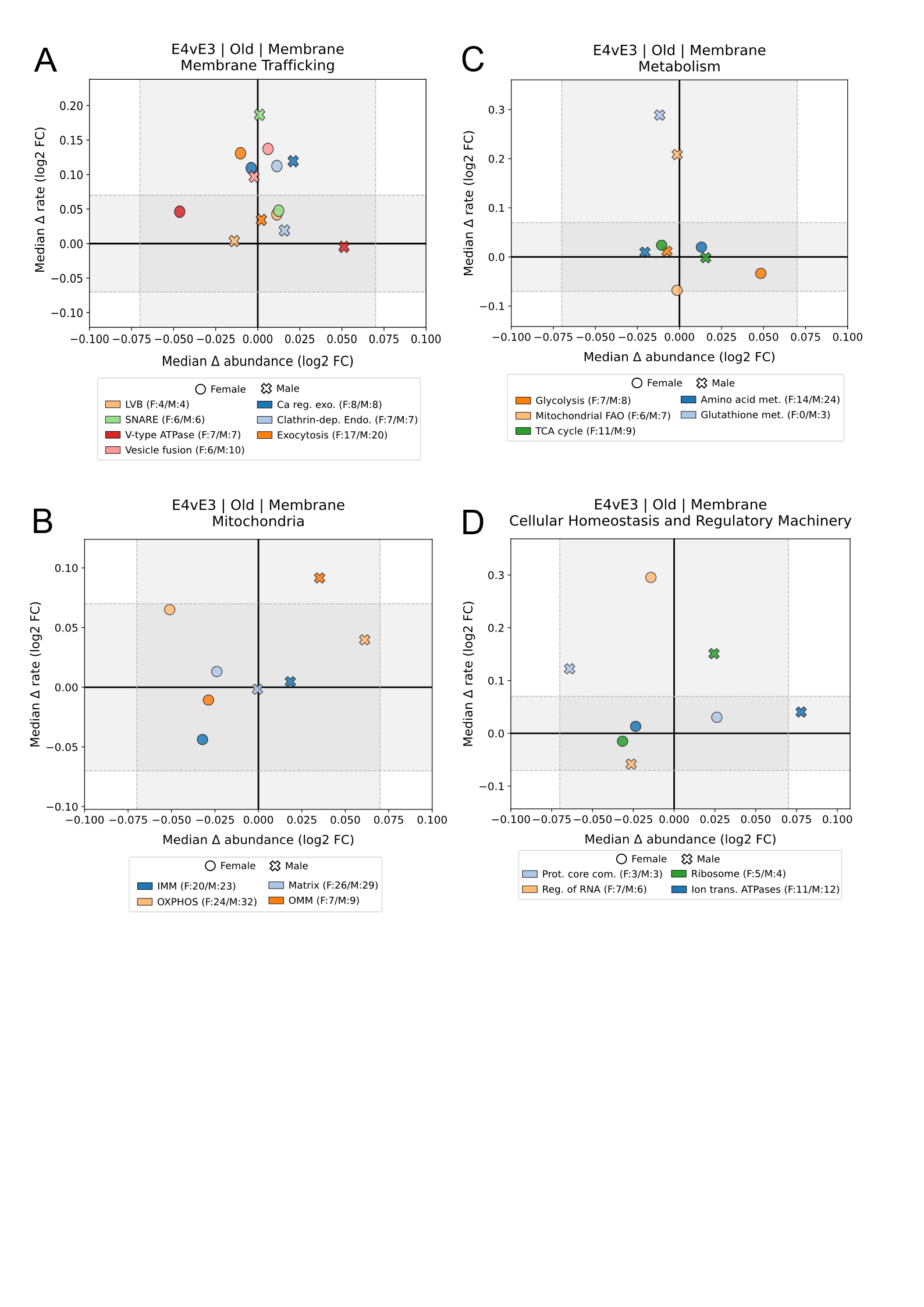

**Supplementary Figure 5. Old E4vE3 Membrane Proteostasis Plots.** Proteostasis plot of normalized protein abundance (x-axis) and rate (y-axis) fold changes of 18-month-old male and female mice organized into functional groups for **(A)** Membrane Trafficking, **(B)** Mitochondria, **(C)** Metabolism, **(D)** and Cellular Homeostasis and Regulatory Machinery ontologies. Fold Change cutoffs for abundance and rate ontologies are |0.07|. Red outlines on boxes represent a significant Wilcoxon p-value of less than 0.05. Ca reg. exo. = Calcium-ion regulated exocytosis, Clathrin-dep. Endo. = Clathrin-dependent endocytosis, LVB = Lysosome Vesicle Biogenesis, Mitophagy = Autophagy of Mitochondrion, IMM = Inner Mitochondrial Membrane, Matrix= Mitochondrial matrix, OMM = Outer Mitochondrial Membrane, OXPHOS= Oxidative Phosphorylation, Amino acid met. = Amino acid metabolic process, Glutathione met. = Glutathione metabolism, Glycolysis = Canonical Glycolysis, Mitochondrial FAO = Mitochondrial fatty acid beta-oxidation, TCA cycle = Tricarboxylic acid cycle, Ion trans. ATPases = Ion transport by P-type ATPases, Prot. Core com. = Proteasome core complex, Prot. Reg. part. = Proteasome regulatory particle, Reg. of RNA = Regulation of RNA splicing.

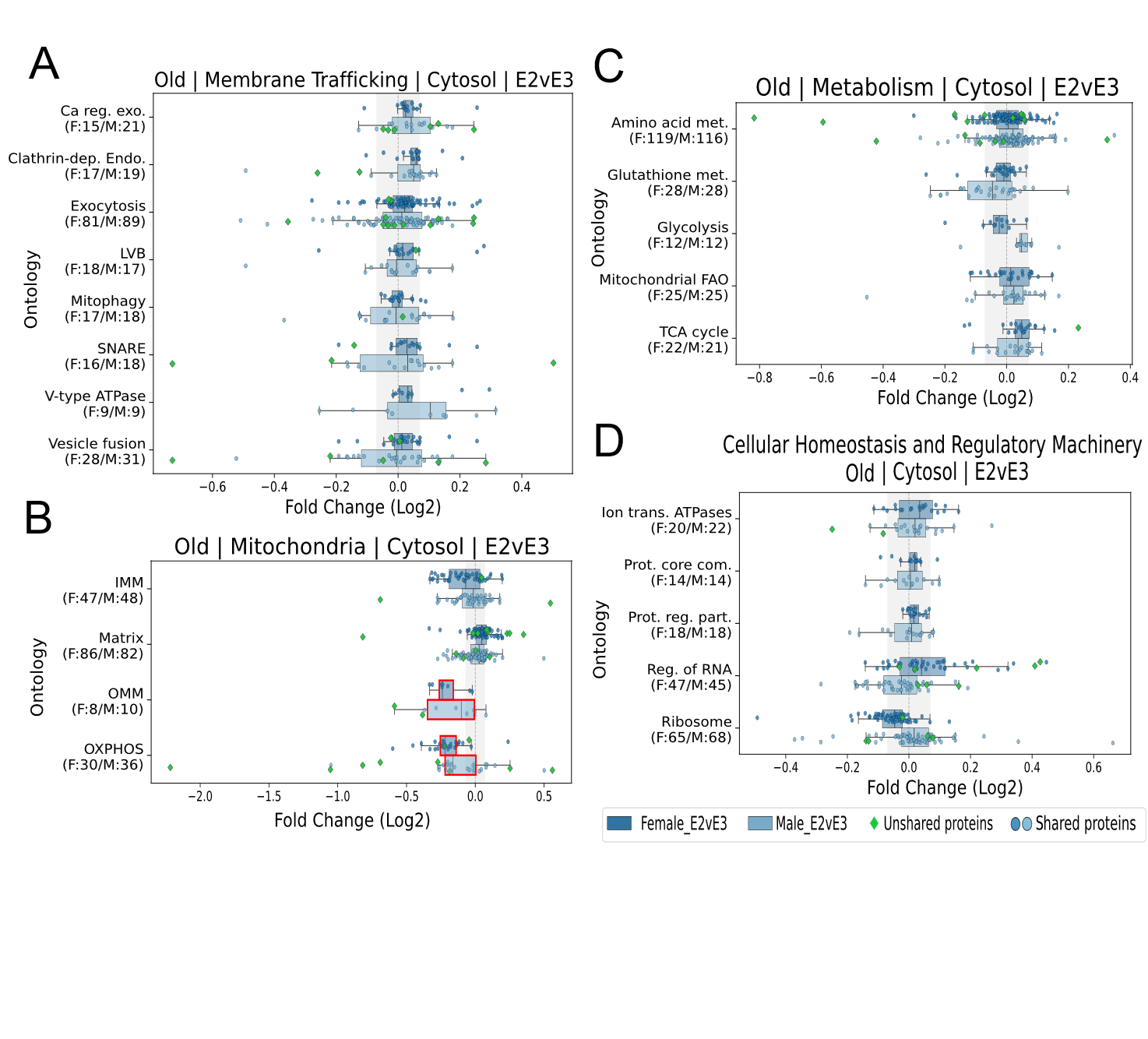

**Supplementary Figure 6**. **Old E2vE3 Cytosolic Boxplots**. Boxplots of normalized protein abundances of 18-month-old male and female mice organized into functional groups for female (top, dark) and male (bottom, light) in **(A)** Membrane Trafficking, **(B)** Mitochondria, **(C)** Metabolism, **(D)** and Cellular Homeostasis and Regulatory Machinery ontologies. X-axis is log2 fold change of ApoE2/ApoE3, y-axis are ontologies. Fold Change cutoffs for abundance and rate ontologies are |0.07|. Red outlines on boxes represent a significant Wilcoxon p-value of less than 0.05. Ca reg. exo. = Calcium-ion regulated exocytosis, Clathrin-dep. Endo. = Clathrin-dependent endocytosis, LVB = Lysosome Vesicle Biogenesis, Mitophagy = Autophagy of Mitochondrion, IMM = Inner Mitochondrial Membrane, Matrix= Mitochondrial matrix, OMM = Outer Mitochondrial Membrane, OXPHOS= Oxidative Phosphorylation, Amino acid met. = Amino acid metabolic process, Glutathione met. = Glutathione metabolism, Glycolysis = Canonical Glycolysis, Mitochondrial FAO = Mitochondrial fatty acid beta-oxidation, TCA cycle = Tricarboxylic acid cycle, Ion trans. ATPases = Ion transport by P-type ATPases, Prot. Core com. = Proteasome core complex, Prot. Reg. part. = Proteasome regulatory particle, Reg. of RNA = Regulation of RNA splicing.

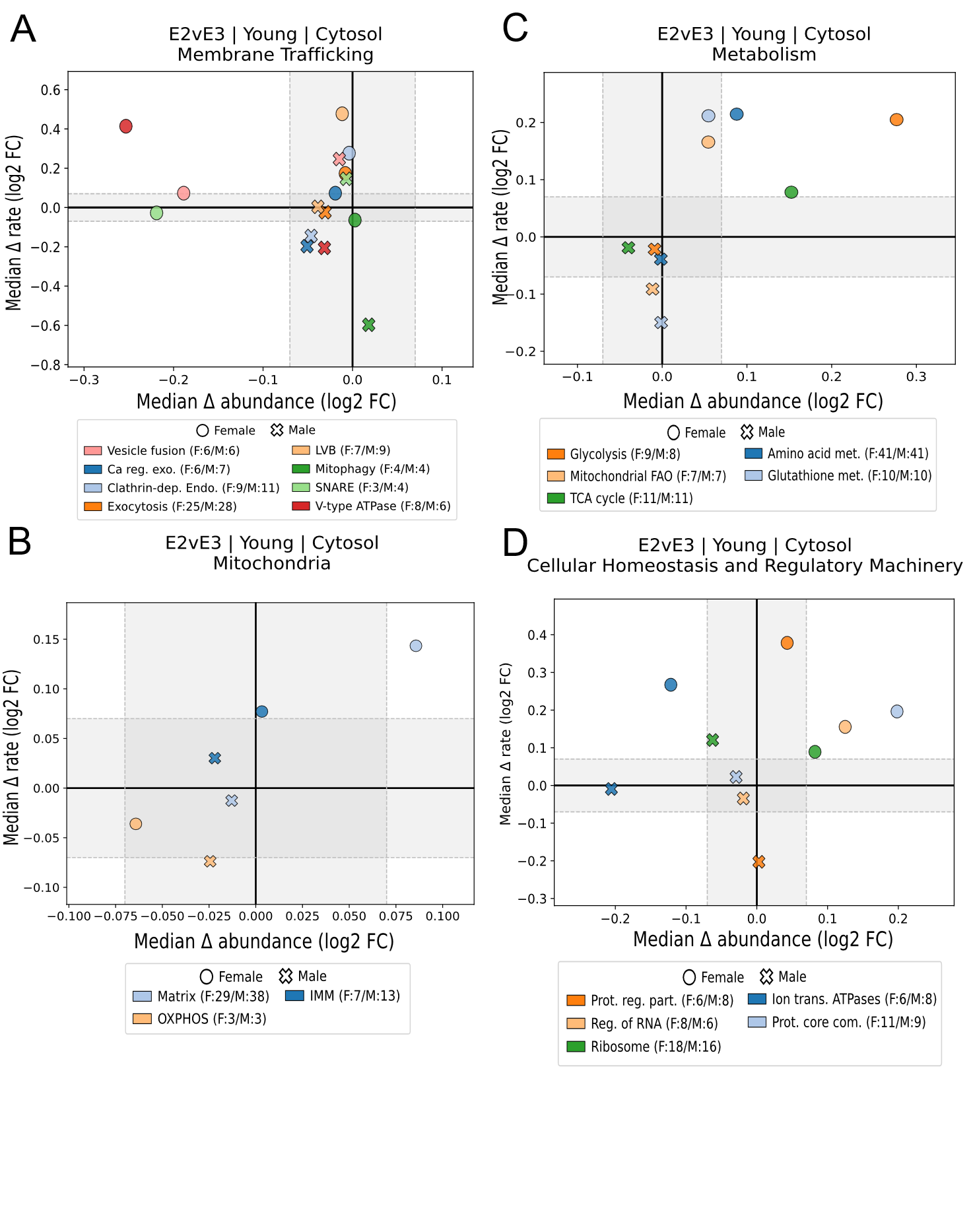

**Supplementary Figure 7. Young E2vE3 Cytosolic Proteostasis Plots**. Proteostasis plots of normalized protein abundance (x-axis) and rate (y-axis) fold changes of 3-month-old male and female mice organized into functional groups for **(A)** Membrane Trafficking, **(B)** Mitochondria, **(C)** Metabolism, **(D)** and Cellular Homeostasis and Regulatory Machinery ontologies. Fold Change cutoffs for abundance and rate ontologies are |0.07|. Red outlines on boxes represent a significant Wilcoxon p-value of less than 0.05. Ca reg. exo. = Calcium-ion regulated exocytosis, Clathrin-dep. Endo. = Clathrin-dependent endocytosis, LVB = Lysosome Vesicle Biogenesis, Mitophagy = Autophagy of Mitochondrion, IMM = Inner Mitochondrial Membrane, Matrix= Mitochondrial matrix, OMM = Outer Mitochondrial Membrane, OXPHOS= Oxidative Phosphorylation, Amino acid met. = Amino acid metabolic process, Glutathione met. = Glutathione metabolism, Glycolysis = Canonical Glycolysis, Mitochondrial FAO = Mitochondrial fatty acid beta-oxidation, TCA cycle = Tricarboxylic acid cycle, Ion trans. ATPases = Ion transport by P-type ATPases, Prot. Core com. = Proteasome core complex, Prot. Reg. part. = Proteasome regulatory particle, Reg. of RNA = Regulation of RNA splicing.

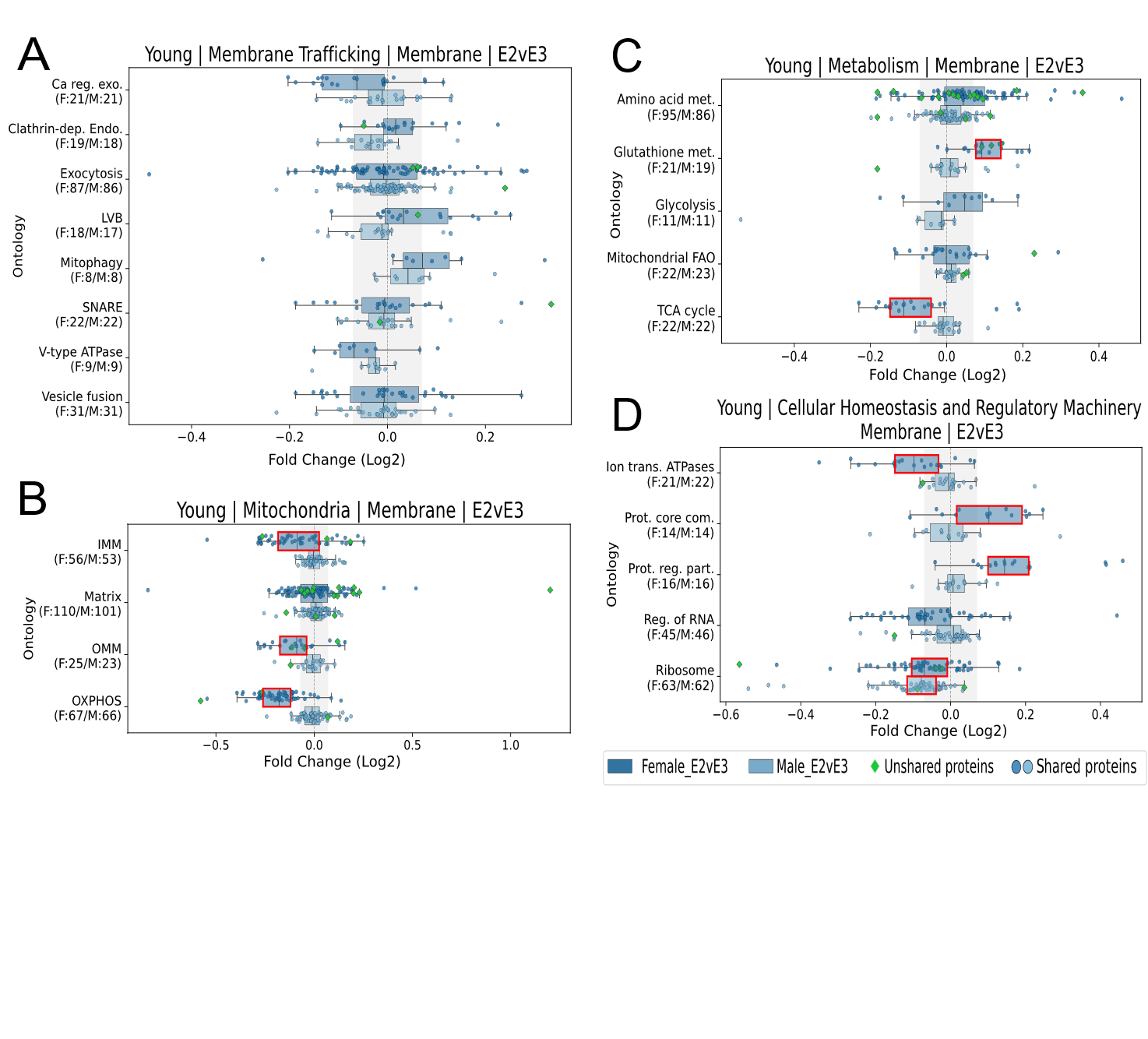

**Supplementary Figure 8. Unique Membrane Ontological Signatures in ApoE2 Females.** Boxplots of normalized protein abundances of 3-month-old mice organized into functional groups for female (top, dark) and male (bottom, light) in **(A)** Membrane Trafficking, **(B)** Mitochondria, **(C)** Metabolism, and **(D)** Cellular Homeostasis and Regulatory Machinery Ontologies. X-axis is log2 fold change of ApoE2/ApoE3, y-axis for are ontologies. Fold Change cutoffs for abundance and rate ontologies are |0.07|. Red outlines on boxes represent a significant Wilcoxon p-value of less than 0.05. Ca reg. exo. = Calcium-ion regulated exocytosis, Clathrin-dep. Endo. = Clathrin-dependent endocytosis, LVB = Lysosome Vesicle Biogenesis, Mitophagy = Autophagy of Mitochondrion, IMM = Inner Mitochondrial Membrane, Matrix= Mitochondrial matrix, OMM = Outer Mitochondrial Membrane, OXPHOS= Oxidative Phosphorylation, Amino acid met. = Amino acid metabolic process, Glutathione met. = Glutathione metabolism, Glycolysis = Canonical Glycolysis, Mitochondrial FAO = Mitochondrial fatty acid beta-oxidation, TCA cycle = Tricarboxylic acid cycle, Ion trans. ATPases = Ion transport by P-type ATPases, Prot. Core com. = Proteasome core complex, Prot. Reg. part. = Proteasome regulatory particle, Reg. of RNA = Regulation of RNA splicing.

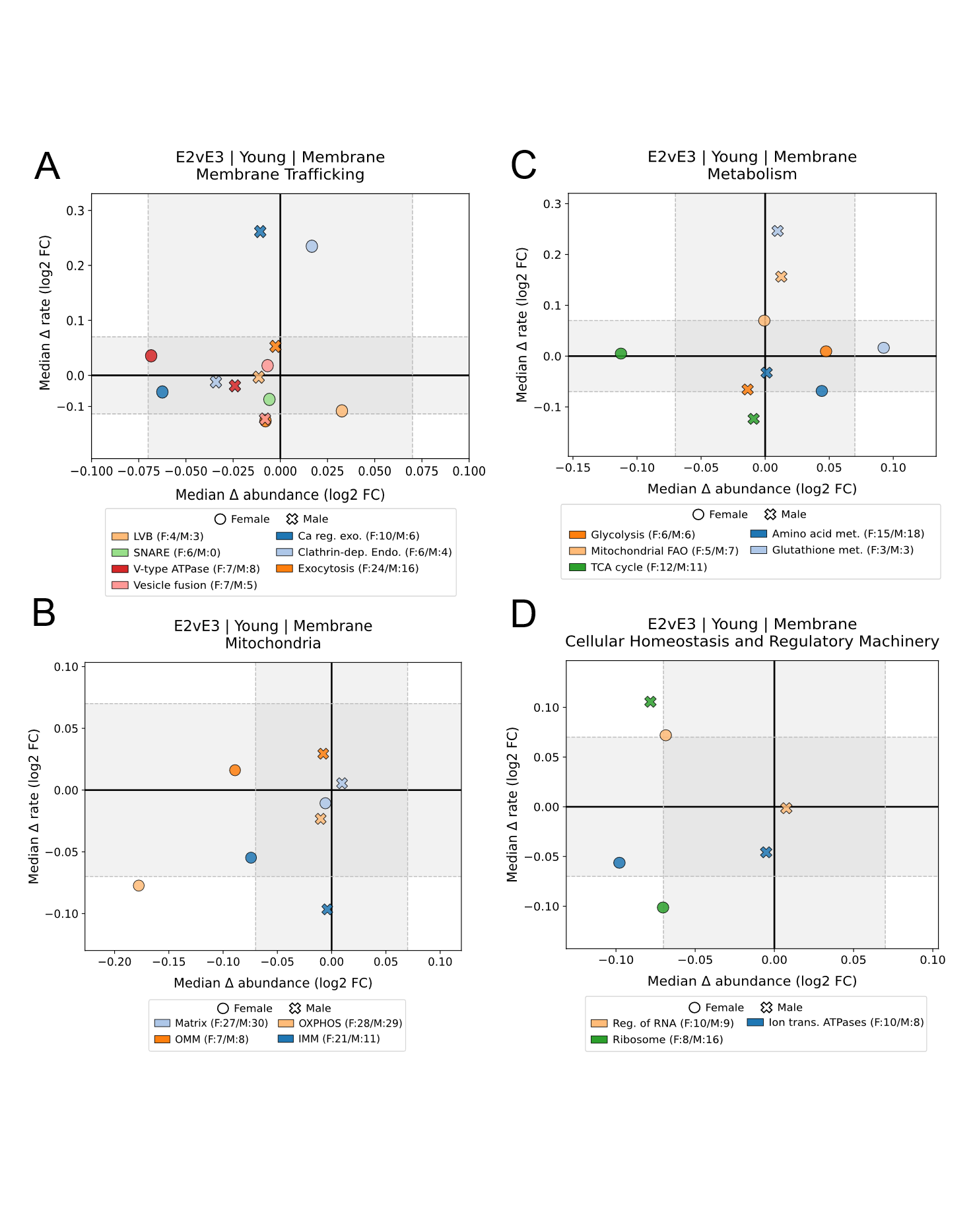

**Supplementary Figure 9. Young E2vE3 Membrane Proteostasis Plots**. Proteostasis plot of normalized protein abundance (x-axis) and rate (y-axis) fold changes of 3-month-old male and female mice organized into functional groups for **(A)** Membrane Trafficking, **(B)** Mitochondria, **(C)** Metabolism, **(D)** and Cellular Homeostasis and Regulatory Machinery ontologies. Fold Change cutoffs for abundance and rate ontologies are |0.07|. Red outlines on boxes represent a significant Wilcoxon p-value of less than 0.05. Ca reg. exo. = Calcium-ion regulated exocytosis, Clathrin-dep. Endo. = Clathrin-dependent endocytosis, LVB = Lysosome Vesicle Biogenesis, Mitophagy = Autophagy of Mitochondrion, IMM = Inner Mitochondrial Membrane, Matrix= Mitochondrial matrix, OMM = Outer Mitochondrial Membrane, OXPHOS= Oxidative Phosphorylation, Amino acid met. = Amino acid metabolic process, Glutathione met. = Glutathione metabolism, Glycolysis = Canonical Glycolysis, Mitochondrial FAO = Mitochondrial fatty acid beta-oxidation, TCA cycle = Tricarboxylic acid cycle, Ion trans. ATPases = Ion transport by P-type ATPases, Prot. Core com. = Proteasome core complex, Prot. Reg. part. = Proteasome regulatory particle, Reg. of RNA = Regulation of RNA splicing.

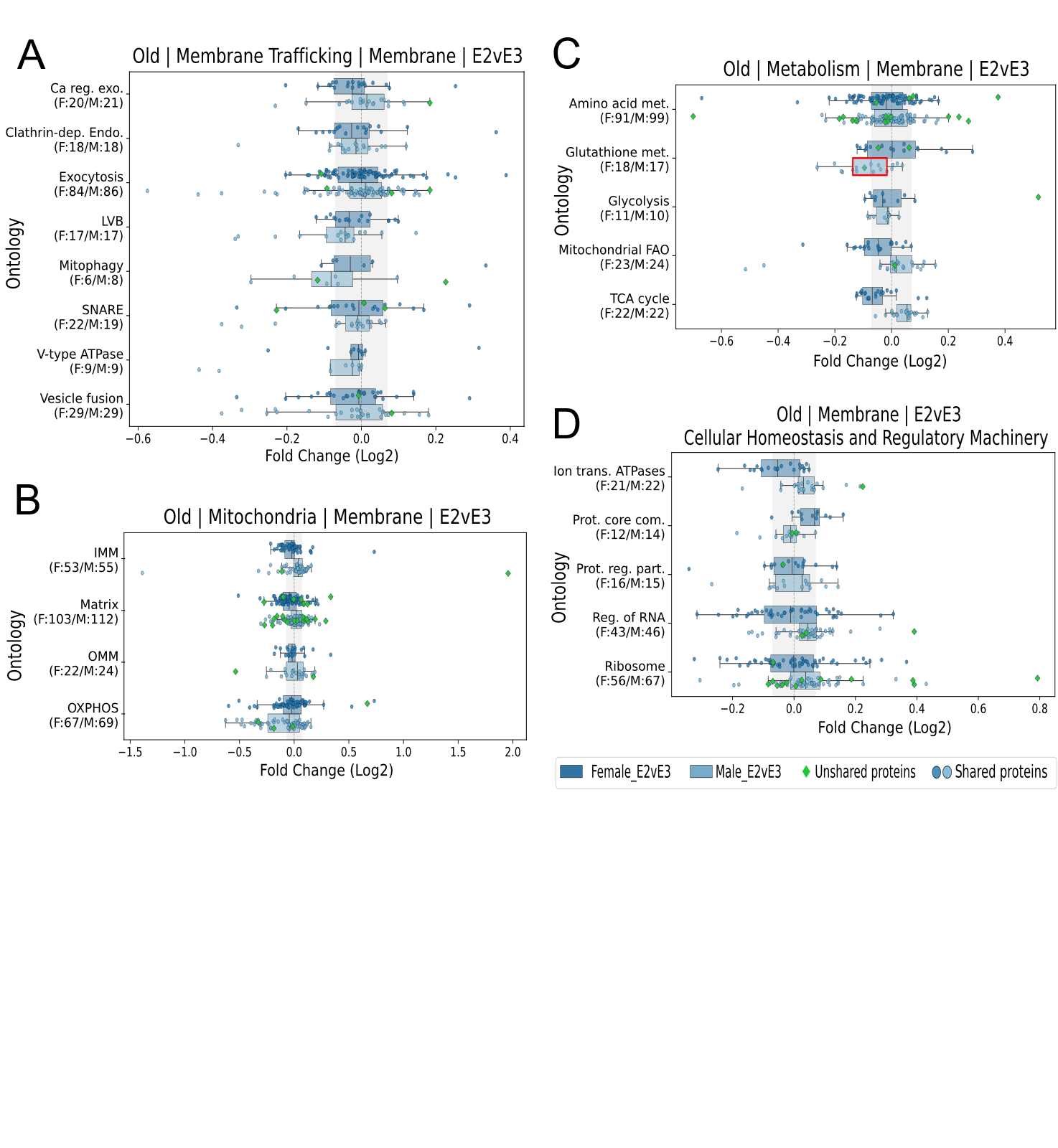

**Supplementary Figure 10. Old E2vE3 Membrane Boxplots**. Boxplots of normalized protein abundances of 18-month-old mice organized into functional groups for female (top, dark) and male (bottom, light) in **(A)** Membrane Trafficking, **(B)** Mitochondria, **(C)** Metabolism, and **(D)** Cellular Homeostasis and Regulatory Machinery Ontologies. X-axis is log2 fold change of ApoE2/ApoE3, y-axis for are ontologies. Fold Change cutoffs for abundance and rate ontologies are |0.07|. Red outlines on boxes represent a significant Wilcoxon p-value of less than 0.05. Ca reg. exo. = Calcium-ion regulated exocytosis, Clathrin-dep. Endo. = Clathrin-dependent endocytosis, LVB = Lysosome Vesicle Biogenesis, Mitophagy = Autophagy of Mitochondrion, IMM = Inner Mitochondrial Membrane, Matrix= Mitochondrial matrix, OMM = Outer Mitochondrial Membrane, OXPHOS= Oxidative Phosphorylation, Amino acid met. = Amino acid metabolic process, Glutathione met. = Glutathione metabolism, Glycolysis = Canonical Glycolysis, Mitochondrial FAO = Mitochondrial fatty acid beta-oxidation, TCA cycle = Tricarboxylic acid cycle, Ion trans. ATPases = Ion transport by P-type ATPases, Prot. Core com. = Proteasome core complex, Prot. Reg. part. = Proteasome regulatory particle, Reg. of RNA = Regulation of RNA splicing.

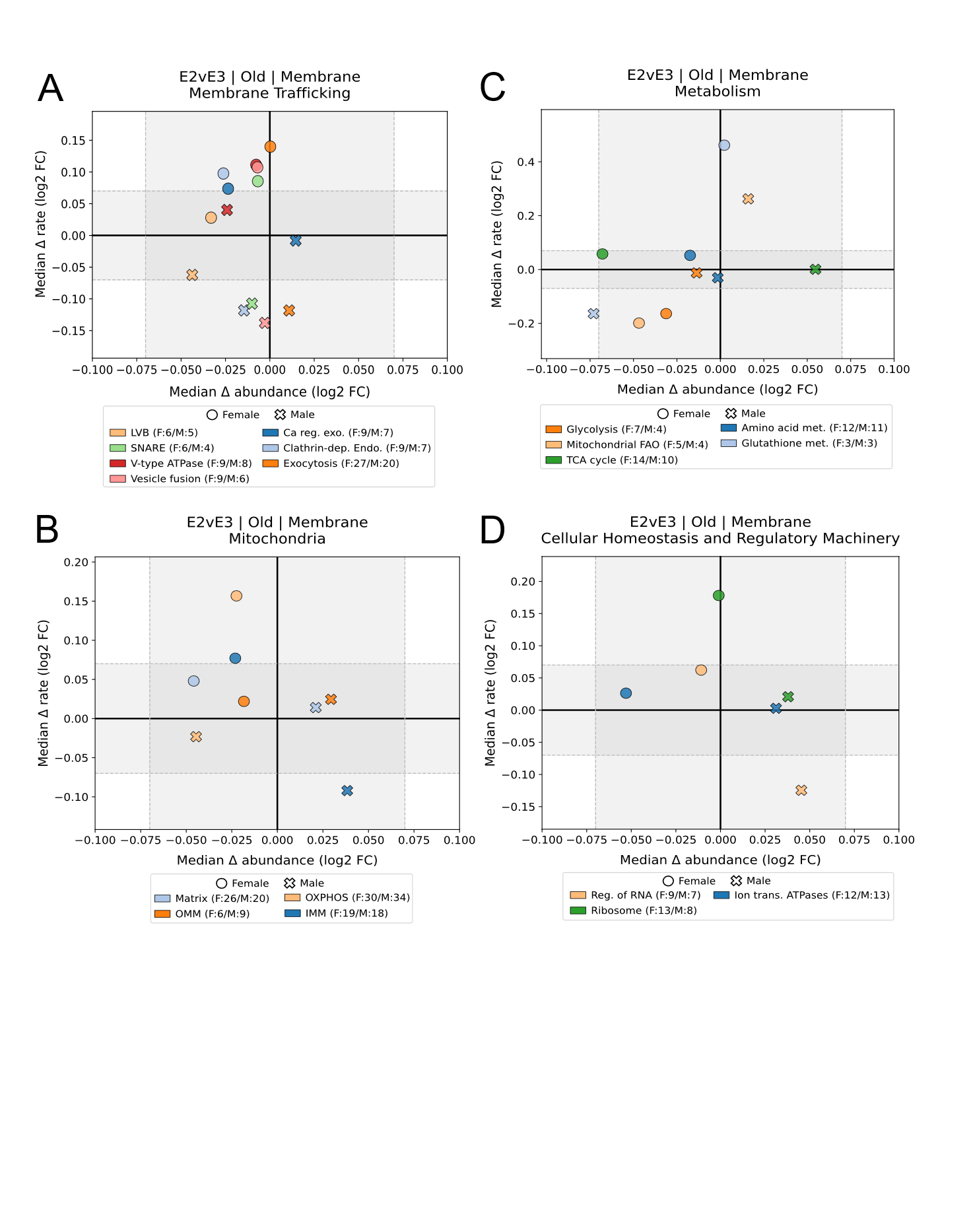

**Supplementary Figure 11. Old E2vE3 Membrane Proteostasis Plots.** Proteostasis plot of normalized protein abundance (x-axis) and rate (y-axis) fold changes of 18-month-old male and female mice organized into functional groups for **(A)** Membrane Trafficking, **(B)** Mitochondria, **(C)** Metabolism, **(D)** and Cellular Homeostasis and Regulatory Machinery ontologies. Fold Change cutoffs for abundance and rate ontologies are |0.07|. Red outlines on boxes represent a significant Wilcoxon p-value of less than 0.05. Ca reg. exo. = Calcium-ion regulated exocytosis, Clathrin-dep. Endo. = Clathrin-dependent endocytosis, LVB = Lysosome Vesicle Biogenesis, Mitophagy = Autophagy of Mitochondrion, IMM = Inner Mitochondrial Membrane, Matrix= Mitochondrial matrix, OMM = Outer Mitochondrial Membrane, OXPHOS= Oxidative Phosphorylation, Amino acid met. = Amino acid metabolic process, Glutathione met. = Glutathione metabolism, Glycolysis = Canonical Glycolysis, Mitochondrial FAO = Mitochondrial fatty acid beta-oxidation, TCA cycle = Tricarboxylic acid cycle, Ion trans. ATPases = Ion transport by P-type ATPases, Prot. Core com. = Proteasome core complex, Prot. Reg. part. = Proteasome regulatory particle, Reg. of RNA = Regulation of RNA splicing.

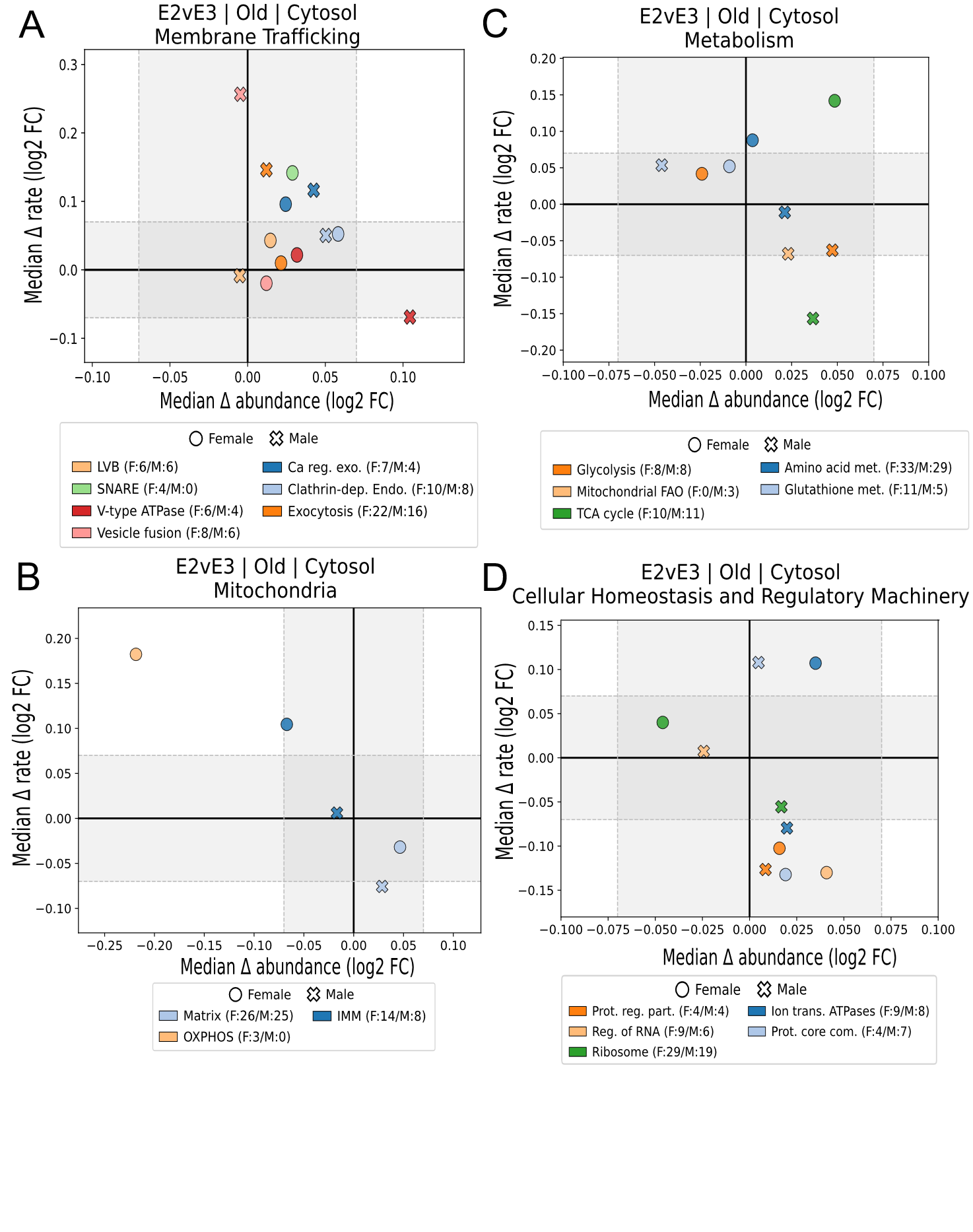

**Supplementary Figure 12. Old E2vE3 Cytosolic Proteostasis Plot.** Proteostasis plot of normalized protein abundance (x-axis) and rate (y-axis) fold changes of 18-month-old male and female mice organized into functional groups for **(A)** Membrane Trafficking, **(B)** Mitochondria, **(C)** Metabolism, **(D)** and Cellular Homeostasis and Regulatory Machinery ontologies. Fold Change cutoffs for abundance and rate ontologies are |0.07|. Red outlines on boxes represent a significant Wilcoxon p-value of less than 0.05. Ca reg. exo. = Calcium-ion regulated exocytosis, Clathrin-dep. Endo. = Clathrin-dependent endocytosis, LVB = Lysosome Vesicle Biogenesis, Mitophagy = Autophagy of Mitochondrion, IMM = Inner Mitochondrial Membrane, Matrix= Mitochondrial matrix, OMM = Outer Mitochondrial Membrane, OXPHOS= Oxidative Phosphorylation, Amino acid met. = Amino acid metabolic process, Glutathione met. = Glutathione metabolism, Glycolysis = Canonical Glycolysis, Mitochondrial FAO = Mitochondrial fatty acid beta-oxidation, TCA cycle = Tricarboxylic acid cycle, Ion trans. ATPases = Ion transport by P-type ATPases, Prot. Core com. = Proteasome core complex, Prot. Reg. part. = Proteasome regulatory particle, Reg. of RNA = Regulation of RNA splicing.

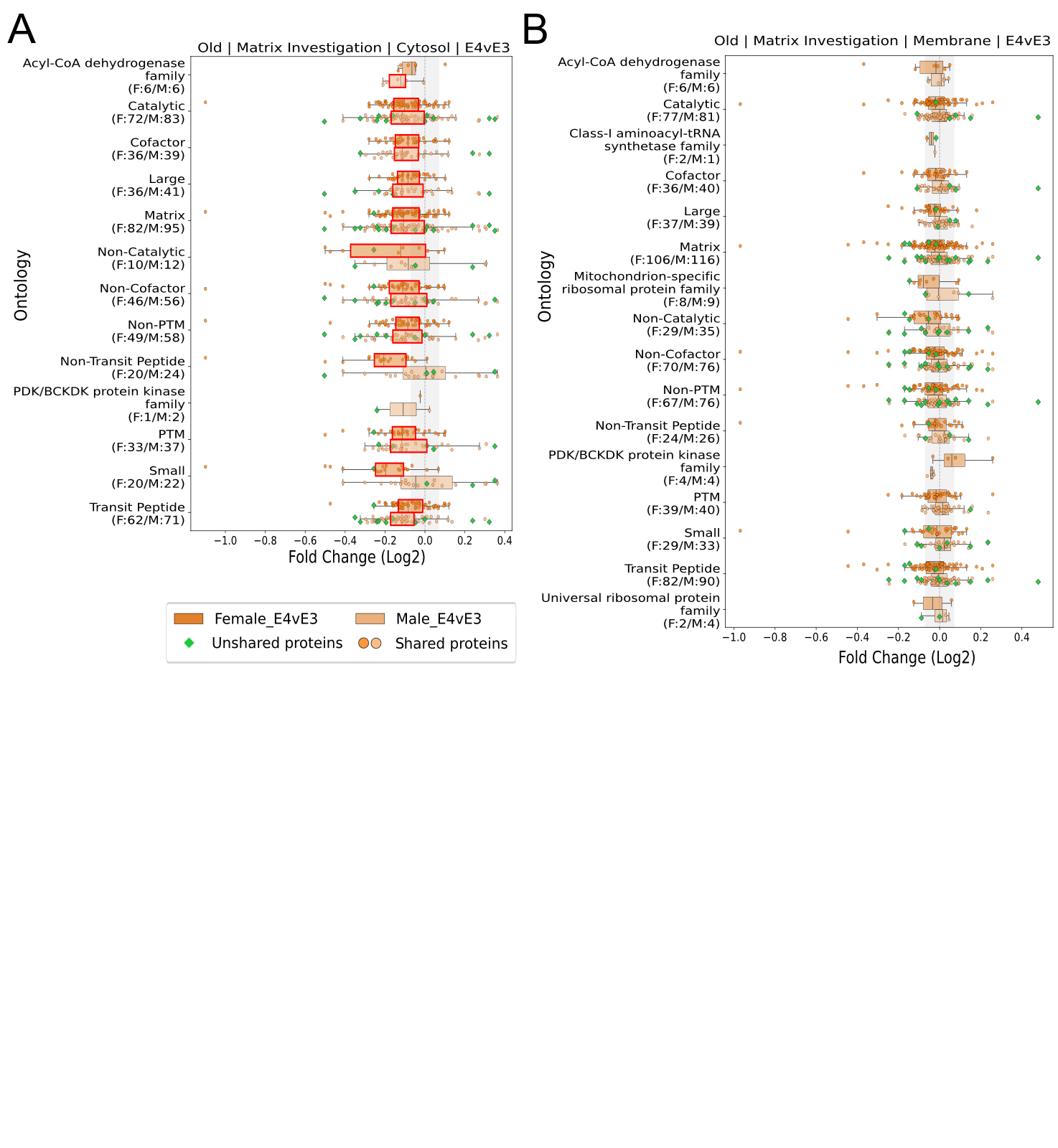

**Supplementary Figure 13.** **Mitochondrial Matrix Investigation.** Boxplots for female (top, dark) and male (bottom, light) for ontology groups created using information from UniProt. Notably, “Non-“ prefixed ontologies are not absolute and simply did not have information for the given grouping at the time.
